# EEG reveals brain network alterations in chronic aphasia during natural speech listening

**DOI:** 10.1101/2023.03.10.532034

**Authors:** Ramtin Mehraram, Jill Kries, Pieter De Clercq, Maaike Vandermosten, Tom Francart

## Abstract

Aphasia is a common consequence of a stroke which affects language processing. In search of an objective biomarker for aphasia, we used EEG to investigate how functional network patterns in the cortex are affected in persons with post-stroke chronic aphasia (PWA) compared to healthy controls (HC) while they are listening to a story.

EEG was recorded from 22 HC and 27 PWA while they listened to a 25-min-long story. Functional connectivity between scalp regions was measured with the weighted phase lag index. The Network-Based Statistics toolbox was used to detect altered network patterns and to investigate correlations with behavioural tests within the aphasia group. Differences in network geometry were assessed by means of graph theory and a targeted node-attack approach. Group-classification accuracy was obtained with a support vector machine classifier.

PWA showed stronger inter-hemispheric connectivity compared to HC in the theta-band (4.5-7 Hz), whilst a weaker subnetwork emerged in the low-gamma band (30.5-49 Hz). Two subnetworks correlated with semantic fluency in PWA respectively in delta- (1-4 Hz) and low-gamma-bands. In the theta-band network, graph alterations in PWA emerged at both local and global level, whilst only local changes were found in the low-gamma-band network. As assessed with the targeted node-attack, PWA exhibit a more scale-free network compared to HC. Network metrics effectively discriminated PWA and HC (AUC = 83%).

Overall, we showed for that EEG-network metrics are effective biomarkers to assess natural speech processing in chronic aphasia. We hypothesize that the detected alterations reflect compensatory mechanisms associated with recovery.

## 1. Introduction

Aphasia is a condition affecting on average 33% and up to 50% post-stroke patients (Gialanella et al., 2010). It is associated with impaired comprehension, reading, writing, and production of language (Flowers et al., 2016; Saur et al., 2006), and due to strong inter-individual variability, the development of efficient diagnostic biomarkers is a matter of interest in language neuroscience.

As reported in functional magnetic resonance imaging (fMRI) studies, aphasia is associated with alteration of physiological neuronal communication between brain regions, suggesting that aphasia is in fact a network disorder (Li et al., 2022). Nevertheless, functional network abnormalities depend on the stage of the pathology. As assessed with fMRI, the (sub-)acute stage is associated with generalized reduced cortical activity and network integration, followed by an increase over the right (non-lesioned) hemisphere and subsequent normalization towards the chronic stage (Hartwigsen & Saur, 2019; Saur et al., 2006; Siegel et al., 2016). The development of the pathology is also reflected in the distribution of connectivity patterns across the cortex. The acute stage of aphasia was shown to feature reduced inter-hemispheric and higher intra-hemispheric connectivity between language and domain-general cortical regions (Siegel et al., 2016). At the chronic stage, activation of right-hemispheric or domain-general regions depends on the severity and properties of the lesion over the left hemisphere, which retains a major role in language recovery, as well as on the difficulty of the task (Brownsett et al., 2014; Kiran & Thompson, 2019; Sims et al., 2016). In contrast, graph property measures showed no differences or higher network integration in persons with aphasia at the acute stage when compared to healthy controls (Chen et al., 2021; Zhu et al., 2017) whilst no evidence exists in the chronic stage.

Electroencephalography (EEG) is a convenient alternative to fMRI for both research and diagnostics due to its reduced cost and higher temporal resolution. To date, it is being extensively used as an informative biomarker for diverse conditions which include epilepsy (Acharya et al., 2013; Taylor et al., 2022), schizophrenia (Canuet et al., 2011; Shin et al., 2011), and dementia (Mehraram et al., 2022; Peraza et al., 2018; Schumacher et al., 2020). Recently it has become also of research interest for the diagnosis of aphasia and prediction of recovery. At both acute and chronic stages, EEG quantitative measures show a shift towards lower frequencies of alpha and beta bands in the spectrum, associated with severity of both cognitive and language impairment (Bentes et al., 2018; Dalton et al., 2021). Notably, task-based event-related potential (ERP) studies found altered cortical responses to vocal stimuli in persons with post-stroke chronic aphasia (PWA) with a loss of inter-hemispheric activity, also associated with the extent of the lesioned region (Behroozmand et al., 2022; Hagoort et al., 1996; Pulvermüller et al., 2004; Spironelli & Angrilli, 2015).

In addition to power-spectrum alterations, EEG is also suitable for investigating cortical network correlates of diverse clinical conditions (Dauwan et al., 2016; Mehraram et al., 2022; Sakkalis, 2011). Network patterns can either be assessed via effective or functional connectivity. The former provides causality information by assessing the mutual predictivity between every couple of signals, and includes biological hypothesis-based methods, e.g. neural mass modelling (Bhattacharya et al., 2011; Moran et al., 2013), and data-driven approaches, e.g. Granger causality (Florin et al., 2010; Soleimani et al., 2022). Functional connectivity is a measure of association between the recorded signals, which can be measured via methods including amplitude correlation and phase-based measures (Vinck et al., 2011). Both approaches are complementary and allow to analyze the same condition from different perspectives (for a review on the topic the reader is referred to Sakkalis (2011)). In fact, EEG network metrics showed diagnostic potential for conditions which include attention-deficit/hyperactivity disorder, depression and dementia (Abbas et al., 2021; Hill et al., 2021; Mehraram et al., 2020).

According to resting-state M/EEG-based graph theory studies, the functional brain connectome in acute aphasia features alterations in its geometrical properties, which include reduced network segregation in delta- and theta-band networks at the acute stage compared to healthy condition (Caliandro et al., 2017), and an association between node degree of Broca area and subsequent language improvement (Nicolo et al., 2015). At the chronic stage, patients show left-hemispheric reduction of connectivity strength in alpha- and beta- bands, with an increase in a local low-gamma band subnetwork compared with healthy people (P. Shah-Basak et al., 2022; Snyder et al., 2021). Furthermore, left-hemispheric alpha and centro-frontal beta connectivity are reportedly associated with alteration of speech processing (P. Shah-Basak et al., 2022).

Despite a certain number of studies involving specific tasks or resting state paradigms, there is a lack of research on the functional network correlates of perception and processing of natural speech in PWA. Given the ecological validity of continuous speech (Brodbeck et al., 2018; Brodbeck & Simon, 2020; Hamilton & Huth, 2018), associated cortical tracking mechanisms and functional network patterns have already been investigated in existing studies on healthy people (Brodbeck & Simon, 2020; di Liberto et al., 2015; Ding & Simon, 2014; Kong et al., 2015; Vanthornhout et al., 2018; Zhang et al., 2019). However, a picture of brain functional network alterations in chronic aphasia is missing.

In a novel manner, we perform an exploratory analysis and use EEG to assess the functional connectome in PWA while they are listening to a naturally spoken story. Beside detecting differential network patterns between groups, we also assess differences in the network architecture via a graph theory analysis, test whether any network pattern is associated with the severity of aphasia, and investigate the diagnostic potential for aphasia of EEG-based network metrics.

## 2. Materials and Methods

### 2.1. Participants

PWA were recruited within the cohort of participants at the stroke unit of the University Hospital Leuven, Belgium (UZ Leuven), within a larger project comprising other research works (Kries et al., 2022). Screening was performed using a Dutch adaptation of the Language Screening Test (LAST) (Flamand-Roze et al., 2011; Schevenels et al., 2020, 2022), and only patients with LAST score equal or below the cut-off score (i.e., 14/15) and with left-hemispheric or bilateral lesion were recruited. For each included patient, the experimental session took place at least six months after the stroke onset (median: 19 months; range: 6-369 months). Healthy control participants (HC) were recruited making sure to match the average age of PWA. The participants had no history of psychiatric or neurodegenerative disorders and gave written consent before participation. The study was approved by the Medical Ethical Committee UZ/KU Leuven (S60007).

Our cohort comprised 22 HC (72 ± 7 years old, 15 males) and 27 PWA (73 ± 11 years old, 20 males). Demographic and behavioral information was assessed through standardized clinical tests as described in detail in the work by Kries and colleagues (2022), and is reported in Table 1. Briefly, the Oxford Cognitive Screen (OCS) (Demeyere et al., 2015) was used to assess participants’ cognition, and language tests comprised the Nederlandse Benoem Test (NBT), i.e. Dutch naming test (van Ewijk et al., 2018), the subtests for semantic and phonological fluency of the Comprehensive Aphasia Test in Dutch (CAT-NL) (Visch-Brink et al., 2014) and the ScreeLing test (El Hachioui et al., 2012). Although seven PWA did not score below the cut-off scores in the behavioral tests on the session day, they had a documented language impairment in the acute phase and were still attending speech-language therapy sessions at the time when the testing sessions took place (Kries et al., 2022).

**Table 1.** Demographic and behavioral data.

|  |  | HC (n = 22) | PWA (n = 27) | Statistics |
| --- | --- | --- | --- | --- |
| Demographics | Age (years old) | 72 ± 7 | 73 ± 11 | $U = 721.5, P = 0.355^a$ |
| | Sex (male/female) | 15/7 | 20/7 | $\chi^2 = 0.206, P = 0.650^b$ |
| Cognition | **OCS total score**<br>Composite %: attention, memory and executive function (Demeyere et al., 2015). | 94.06 ± 4.69 | 77.17 ± 15.68 | $U = 469, P < 0.001^a$ |
| Language tests | **NBT total score**<br>Participants were asked to describe a series of images in one word each (van Ewijk et al., 2018). | 271.00 ± 4.16 | 228.58 ± 57.75 | $U = 408.5, P < 0.001^a$ |
| | **CAT-NL semantic**<br>(n. of words)<br>Within cognitive subtests of CAT-NL (Visch-Brink et al., 2014). Participants were asked to pronounce words within a certain semantic category (animals) within one minute. | 22.32 ± 6.30 | 12.85 ± 7.09 | $U = 468, P < 0.001^a$ |
| | **CAT-NL phonologic**<br>(n. of words)<br>Within cognitive subtests of CAT-NL (Visch-Brink et al., 2014). Participants were asked to pronounce words starting with a certain letter within one minute. | 13.14 ± 5.18 | 8.44 ± 6.07 | $U = 547, P = 0.010^a$ |
| | **ScreeLing total score**<br>Three parts with each four different tasks, covering semantics, phonology and syntax (El Hachoui et al., 2012). | 69.86 ± 2.51 | 62.11 ± 9.43 | $U = 479, P < 0.001^a$ |

<sup>a</sup>Unpaired Mann–Whitney U-test<sup>b</sup>Chi-squared test

### 2.2. EEG experiment

Participants listened to a 25-minute long fairy tale titled “De Wilde Zwanen” (“The Wild Swans”) by Hans Christian Andersen, narrated by a female Flemish-native speaker. To ensure participants’ attention to the stimulus, a break was introduced every five minutes and questions about the story to that point were asked. The speech stimulus was presented binaurally through ER-3A insert phones (Etymotic Research Inc, IL, USA) using the software APEX (Francart et al., 2008). For each participant stimulus intensity was set at 60 dB SPL plus half of the pure tone average (PTA) of the individual audiometric thresholds at 250, 500 and 1000 Hz, to ensure audibility. All sessions took place in a soundproof room with Faraday cage at the Department of Neurosciences, KU Leuven. EEG signals were obtained while participants listened to the natural speech using a Biosemi ActiveTwo (Amsterdam, Netherlands) high-density cap with 64 Ag/AgCl electrodes distributed according to the 10-20 system, as shown in Figure 1 (Oostenveld & Praamstra, 2001). Recordings were sampled at 8192 Hz.

**Figure 1.**
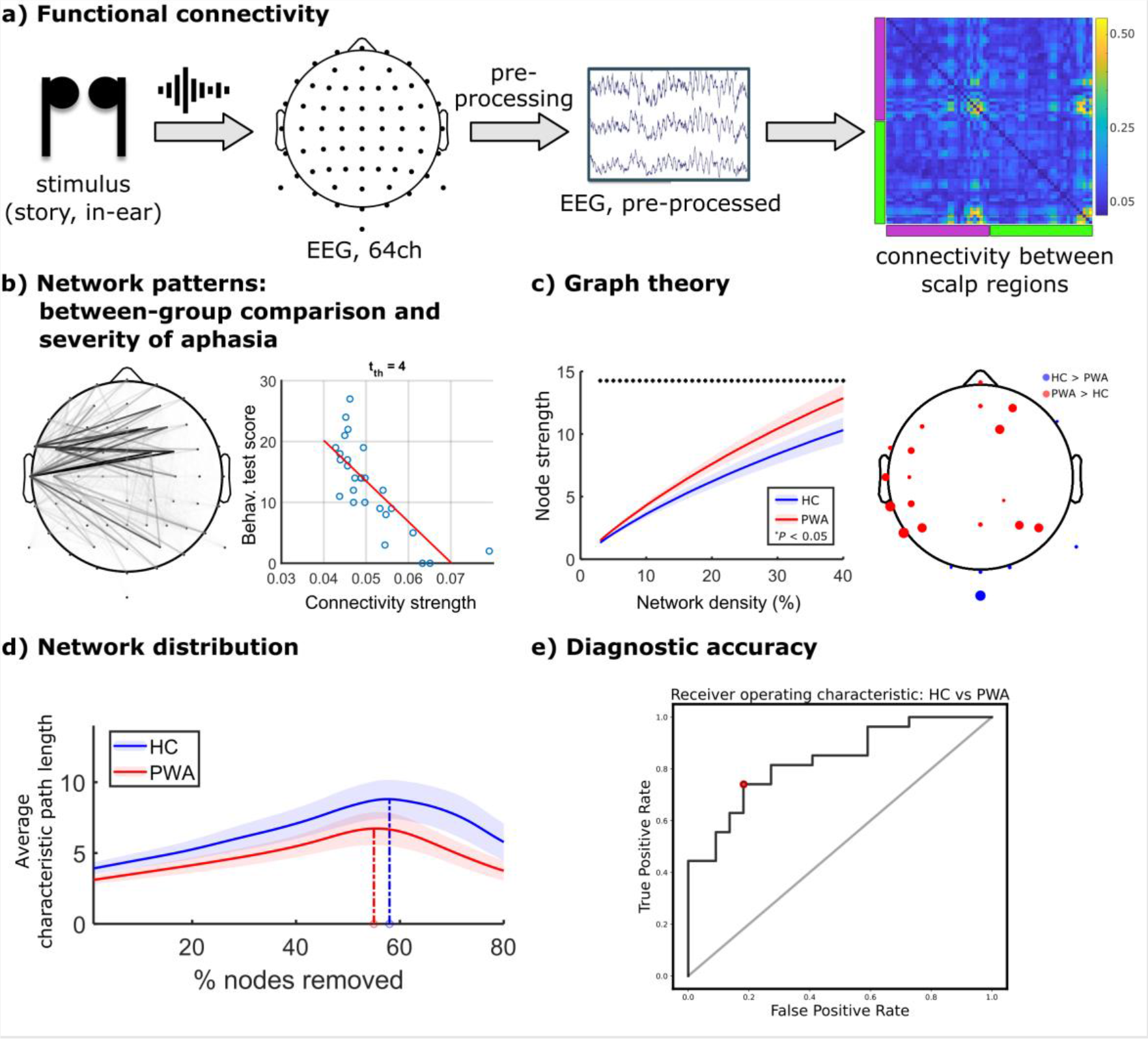
Methodological workflow. a) EEG is recorded while the participants listen to a story, and functional connectivity is extracted from the recorded signals. On the connectivity matrix, purple bar: left hemisphere; green bar: right hemisphere. b) Group differences and correlations with behavioural test scores were assessed. c) Graph measures were computed and compared between groups within the frequency bands with significantly different network patterns. Blue: HC; red: PWA. On the topography, blue: HC > PWA, red: PWA > HC. d) Targeted node-attack to investigate the network distribution in both groups. Blue: HC; red: PWA. e) Diagnostic accuracy analysis with implementation of a support vector machine classifier .

### 2.3. EEG data pre-processing

Pre-processing of the EEG data was automated and was implemented through in-house routines and the Automagic toolbox (Pedroni et al., 2019) v.2.6 on MATLAB v.9.11 (The MathWorks Inc., Natick, MA, USA, 2021). Signals were high-pass filtered with a second-order zero-delay Butterworth filter with cut- off frequency at 0.1 Hz, referenced to Cz, and line noise (50 Hz) was removed using the ZapLine method (de Cheveigné, 2020). The recordings were subsampled to 512 Hz, and channels with correlation to their random sample consensus lower than 80% were deemed bad and removed through the EEGLAB plugin clean_rawdata() (http://sccn.ucsd.edu/wiki/Plugin_list_process) (average number of removed channels: 2 ± 2). Artifactual segments of data were also replaced with a temporal interpolation using the Artifact Subspace Reconstruction (ASR) (Mullen et al., 2015). The resulting data underwent an independent component analysis (ICA), and classification of the components was automatically performed through the EEGLAB plugin ICLabel (Pion-Tonachini et al., 2019), where only components classified as “brain” or “other” (i.e. mixed components) with probability higher than 50% were preserved (average number of removed components: 26 ± 7). The obtained data were projected back to the channel space, and visually inspected as a further quality-check. Before any further analysis, all signals were average-referenced.

### 2.4. Weighted phase lag index

Functional connectivity between scalp EEG signals was measured with the weighted phase lag index (WPLI) (Vinck et al., 2011), mathematically defined as:

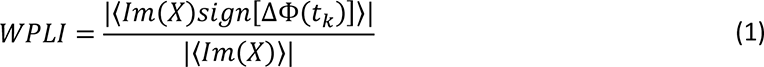

where *X* is the cross-spectrum between the signals, *Im(X)* is its imaginary part, Δφ is the phase difference between the signals, t_k_ is the time step with k = 1, 2, …, *N* and *sign* is the signum function. WPLI values range between 0 and 1, and are insensitive to volume conduction (Peraza et al., 2012), i.e., tend to zero when signals have almost-zero- or zero-lagging synchronization as commonly occurring between conducting signals. WPLI was obtained as implemented in the Fieldtrip toolbox (Oostenveld et al., 2011); EEG signals were first split in three-seconds segments with 50% overlap, then were transformed to the frequency domain using Windowed Fourier Transform (Hanning taper, 0.5 Hz frequency step), and WPLI was computed and averaged across frequency bins. Connectivity was measured within the delta- (1-4 Hz), theta- (4.5-7 Hz), alpha- (7.5-14.5 Hz), beta- (15-30 Hz), low- gamma- (30.5-49 Hz) and mid-gamma-band (50-90 Hz).

### 2.5. Graph theory

Graph theory analysis was performed using functions implemented in the Brain Connectivity Toolbox (BCT) for MATLAB (Rubinov & Sporns, 2010). To compute the network measures, we applied proportional thresholding on the graphs and set to zero the weakest connections. Threshold values (PT%), or network densities, spanned between 3-40% of the total number of edges, and the connectivity weights were preserved in the thresholded graphs in order to reduce the dependence of graph measures on the network density (Bohr et al., 2013; Kaiser, 2011; Langer et al., 2013; Mehraram et al., 2020; Peraza et al., 2015; Rubinov et al., 2009). To prevent any group bias associated with functional connectivity strength, all connectivity matrices were normalized by their maximum weight before computing the network metrics (Onnela et al., 2005). The computed weighted measures comprised: node strength, i.e. total weights of the connections to the node; clustering coefficient, i.e. the extent to which node’s neighbors are connected between each other; eccentricity, i.e. the longest path between the node and all other nodes; characteristic path length, i.e. average shortest path between all pair of nodes; small-worldness, i.e. the ratio between normalized clustering coefficient and normalized characteristic path length; modularity, i.e. the extent to which a network is segregated. Graph measures were computed at each PT%. Further details on the graph theory metrics are reported in Supplementary materials.

### 2.6. Network distribution

To investigate the type of network architecture, we used a targeted node-attack approach as it was described in previous works (Barabasi & Albert, 1999; Kaiser et al., 2007; Mehraram et al., 2020; Stam et al., 2009). For each unthresholded graph, we iteratively removed the nodes with the highest strength, computed the characteristic path length, and obtained its trend with respect to the percentage of removed nodes. An earlier peak is associated with a more scale-free-like as compared to a random or small-world architecture. In order to assess the type of network distribution, we extracted for each subject the percentage of removed nodes at which a peak in the characteristic path length occurred.

### 2.7. Statistical analysis

#### Differential network patterns

EEG network topographies were compared between PWA and HC group using the Network Based Statistics (NBS) toolbox (Zalesky et al., 2010). A test statistics threshold (*tth*) was chosen, graph edges were F-tested against the null-hypothesis of equal average connectivity between groups, and only connections with supra-threshold t-values were preserved. Network components were then built as clusters across the surviving edges, their sizes were measured, and data were permuted between groups. These steps were iteratively performed 5000 times, and the largest component sizes were measured. The final output of the NBS is a family wise error rate-corrected *P*-value for each network component, obtained as a ratio between number of iterations at which the largest component was equally or greater sized compared to the current component and the total number of iterations. We performed the NBS within a range of *tth* values (Zalesky et al., 2010) between one and thirty and visualized the corresponding graphs when the outcomes were significant (*P* < 0.05). To infer the direction of significance, we visualized the distribution within the two groups of the average strength of the NBS connections (WPLI_NBS_). Difference between groups was tested at all frequency bands. Given the exploratory purpose of our analysis, results were not corrected for number of frequency bands.

#### Aphasia severity

We also used the NBS to test whether any EEG subnetwork was associated with aphasia severity. To this purpose, the behavioral scores were included as covariates of interest and an F-test was performed to test whether any correlating network component in the PWA group exists. If any correlation was found, the direction of significance was assessed by visualizing the distribution of the clinical score with respect to WPLI_NBS_.

#### Graph theory

Graph measures were obtained within the frequency ranges for which the NBS analysis yielded a significant outcome and were compared between groups at each network density (PT%). Nodal measures, i.e., node strength and clustering coefficient, were locally tested for each node (64 electrodes, uncorrected for multiple tests) as well as globally on average across nodes with Mann-Whitney U-tests (*P* < 0.05).

#### Network distribution

To compare the distribution of connections within the network, we extracted the percentage of nodes at which a peak in the trend of the average characteristic path length occurred and compared the values between HC and PWA with a Mann-Whitney U-test (*P* < 0.05).

### 2.8. Support vector machine

We investigated the diagnostic potential of WPLI_NBS_ and graph measures within the frequency bands which yielded any significant outcome from the statistical analysis. We implemented a linear support vector machine (SVM) classifier using the Scikit-Learn (v. 0.24.2) library in Python (Pedregosa et al., 2011) to test the accuracy of classification in PWA and HC groups. We implemented a leave-one-out cross-validation, and at each training step the regularization parameter was chosen within a range spanning from 10^-6^ to 10_6_ by means of a cross-validation approach within the training set (5-fold). Given the known effect of aging on brain functional connectivity (Damoiseaux, 2017; Geerligs et al., 2015) age was included as feature in the classifier. For each subject, the WPLI_NBS_ and graph metrics were obtained from the rest of the cohort and were used to train the classifier. We then obtained a mean across the tested subjects of the ranking of the weights assigned to the variables, accuracy, optimal working point as best tradeoff between the rate of true and false detected positives (Fluss et al., 2005; Perkins & Schisterman, 2005), and area under the receiver operating characteristic (AUROC) curve.

## 3. Results

### 3.1. Demographic data

The two groups did not show any significant difference in age (Mann-Whitney U-test, *U* = 721.5, *P* = 0.355), and had a comparable sex distribution (Chi-squared test*, χ*^2^ = 0.206, *P* = 0.650). The HC group performed better in the cognition task (*U* = 469, *P* < 0.001), NBT (*U* = 408.5, *P* < 0.001), CAT-NL (*U* = 547, *P* = 0.010) and ScreeLing (*U* = 479, *P* < 0.001).

### 3.2. Network patterns: between-group comparison

We used the NBS to test whether network patterns differed between PWA and HC. The outcome of this analysis is shown in Figure 2A and reported in Table 2. We found a stronger network component in PWA within the theta-band (5 < *t*_*th*_ < 20, *P* < 0.007, uncorrected for six frequency bands), which comprised connections between the left temporal scalp region and right frontal and parietal nodes. A significant network component also emerged in the low-gamma-band (2 < *t*_*th*_ < 12, *P* < 0.040, uncorrected for six frequency bands), featuring weaker connectivity in PWA within the left-parietal nodes, between occipital and frontal areas, and between left-parietal and right-temporal nodes. No significant components were found for the other frequency bands.

**Figure 2.**
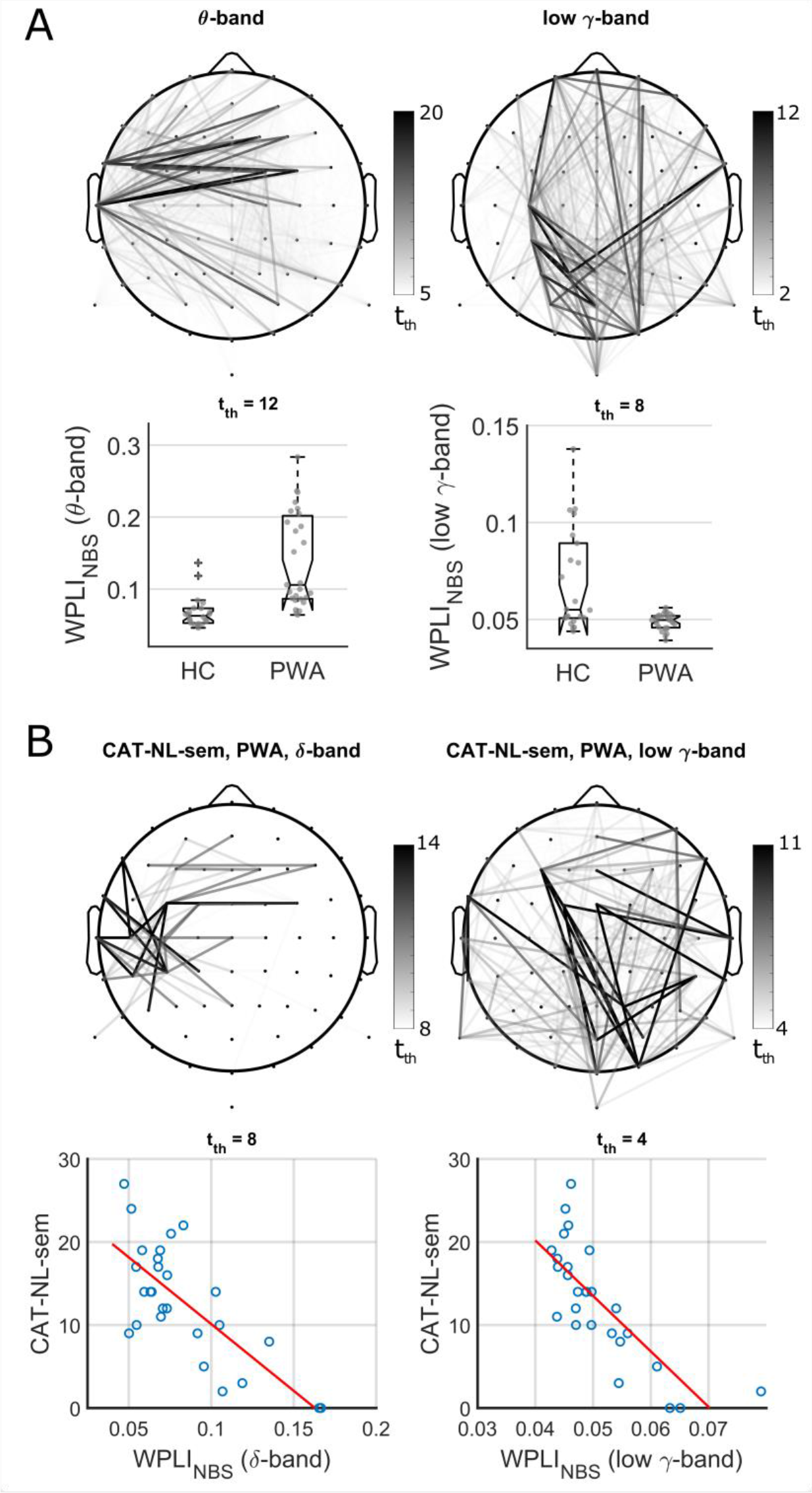
Results of the NBS analysis. Darker edges in the topographies represent connections surviving higher *tth* values. (A) Differential network components between groups were found within the theta (θ)-band and the low-gamma (γ)-band. The boxplots show by way of example the group WPLINBS distributions for the θ- and low- γ-band respectively at *t*_*th*_ = 12 (*P* < 0.001) and *t*_*th*_ = 8 (*P* = 0.011), and distributions at the other *tth* values are reported in Supplementary materials. (B) A significant negative correlation emerged between the CAT-NL semantic fluency score and network components in the delta (δ)- and low-γ-bands. The scatter plots were reported respectively for *t*_*th*_ = 8 (*P* = 0.024) and *t*_*th*_ = 4 (*P* = 0.014), and distributions for the other *t*_*th*_ values are reported in Supplementary materials.

**Table 2.** Significant differences in network metrics between PWA and HC.

| NBS |  |  |  |  |
| --- | --- | --- | --- | --- |
| Significant components: theta- and low-gamma-bands | theta-band |  | low-gamma-band |  |
|  | PWA > HC |  | PWA < HC |  |
| Graph theory |  |  |  |  |
| Graph measure | Local | Global | Local | Global |
| Node strength | • PWA > HC: left temporal, frontal, right parietal regions<br>• PWA < HC: occipital region | PWA > HC | • PWA > HC: left temporal, central regions<br>• PWA < HC: parietal region | No significant differences |
| Clustering coefficient | • PWA > HC: posterior left, central and and anterior right regions | PWA > HC | • PWA > HC: left and right temporal regions | No significant differences |
| Eccentricity | • PWA < HC: all nodes | PWA < HC | • PWA < HC: widespread | PWA < HC |
| Average characteristic path length | X | PWA < HC | X | No significant differences |
| Small-worldness | X | PWA < HC | X | No significant differences |
| Modularity | X | No significant differences | X | No significant differences |

### 3.3. Network patterns: aphasia severity

We explored whether severity of aphasia was associated with any functional subnetwork. To this purpose, we used the NBS and performed F-tests within the PWA group including each behavioral score as co-variate of interest for all frequency bands. We detected two components respectively within the delta-band- (8 < *t*_*th*_ < 14, *P* < 0.038, uncorrected for six frequency bands) and the low- gamma-band-networks (4 < *t*_*th*_ < 11, *P* < 0.035, uncorrected for six frequency bands) that negatively correlated with the semantic fluency score of the CAT-NL, as shown in Figure 2B. No significant correlation was found for the other clinical scores.

### 3.4. Graph theory

Weighted graphs were thresholded from 3 to 40% network densities and graph measures were computed at each density within the theta- and low-gamma-band networks, which yielded significant between-group differences from the NBS. The obtained metrics were compared between groups both locally and globally, as shown in Figure 3 and reported in Table 2. In the theta-band, PWA showed higher average node strength and clustering coefficient, and lower eccentricity, characteristic path length and small-worldness. Although between-group differences in node strength, eccentricity and small-worldness were consistent across PT%, a difference in clustering coefficient emerged only for the highest density values (two-tailed Mann-Whitney U-tests, *P* < 0.05). The node-level analysis revealed that the higher strength in PWA was driven by the left temporal and right frontal and parietal nodes, whilst most clustered regions in PWA comprised the left-occipital and right-frontal nodes. All regions showed consistently higher eccentricity in HC compared to PWA. Although the node strength in PWA was higher on average, lower values compared to HC were found locally for nodes within the occipital scalp region (two-tailed Mann-Whitney U-tests, *P* < 0.05, uncorrected for 64 nodes). In the low-gamma-band-network none or only for few PT% values a between-group differences was detected for the average measures (Figure 3B). Node strength, characteristic path length and small-worldness were not different between groups, whilst the clustering coefficient was higher in PWA for lower values of PT% and the eccentricity was lower in PWA for the highest graph densities (two-tailed Mann-Whitney U-tests, *P* < 0.05). Nevertheless, local differences emerged for all nodal measures (two-tailed Mann-Whitney U-tests, *P* < 0.05). Nodes over the left temporal regions were more strongly connected and clustered in PWA. In addition, one node over the right parietal region and the whole right superior temporal region showed respectively higher strength and clustering coefficient. Similarly to the theta-band network, the differences in eccentricity were widespread across all nodes, although less consistent across PT%. Modularity was not different between groups in either frequency bands.

**Figure 3.**
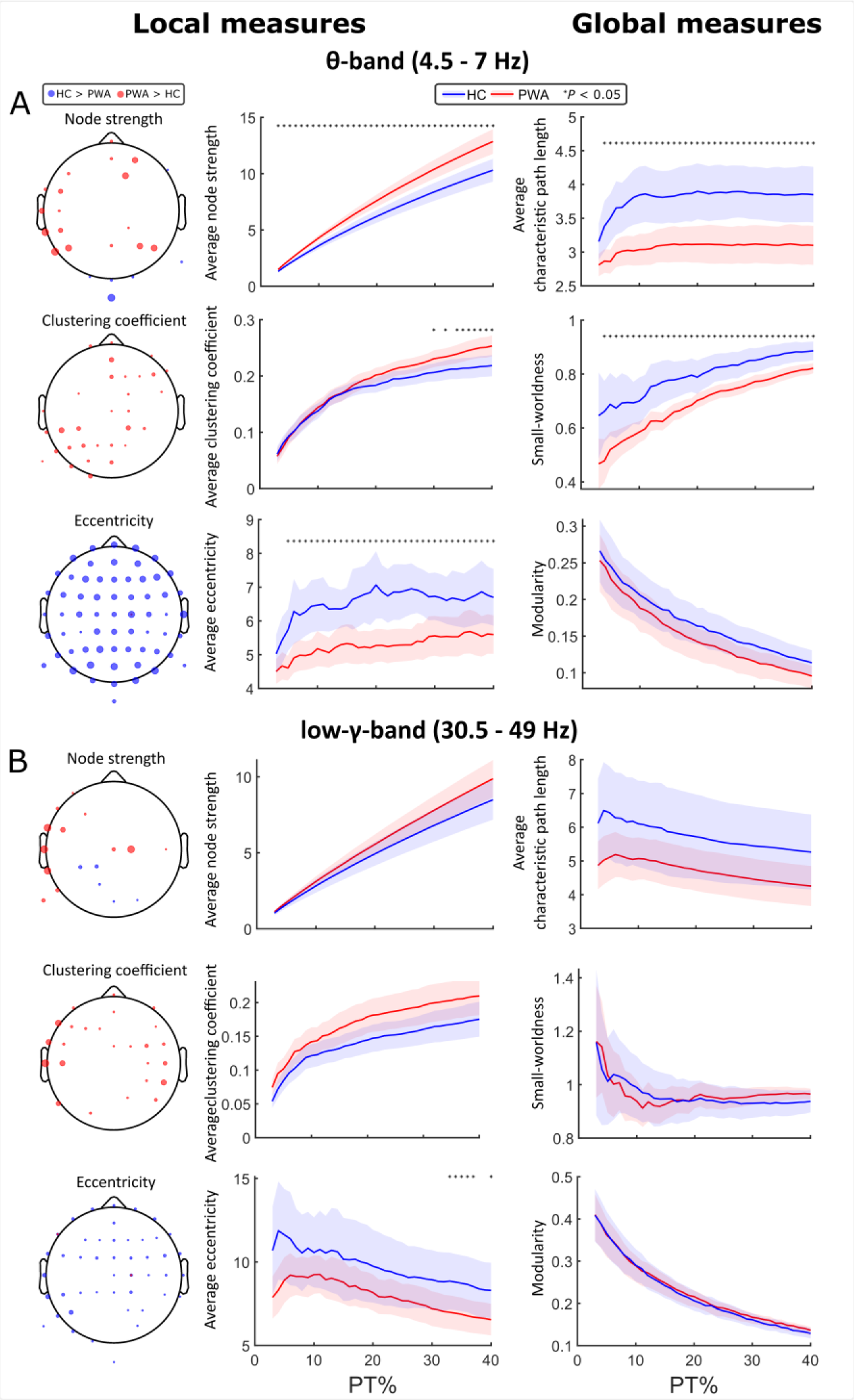
Results of the graph theory analysis. Measures are compared between groups on average and locally with respect to the network density. Blue line: HC group; red line: PWA group. Dotted lines: 95% confidence interval. Blue node: graph measure higher in HC compared with PWA; red node: graph measure higher in PWA compared with HC. Size of the node is proportional to the occurrence of test-significance across PT% values. Statistics are uncorrected for number of nodes. (A) Outcome for the theta (θ)-band network. All measures were different between groups except for the modularity; local measures showed node-specific between-group differences. (B) Outcome for the low-gamma (γ)-band network. Inconsistent differences emerged globally, whilst we found consistent local differences.

### 3.5. Network distribution

To further investigate the network architecture, we performed a targeted node-attack and extracted the trend of the average characteristic path length with respect to the percentage of removed nodes. For the theta-band network, the Mann-Whitney U-test yielded a result close to significance for an earlier peak in the PWA group compared to the HC (*U* = 640.5, *P* = 0.07). No difference was found within the low-gamma-band network. The outcome of this analysis is reported in Figure 4.

**Figure 4.**
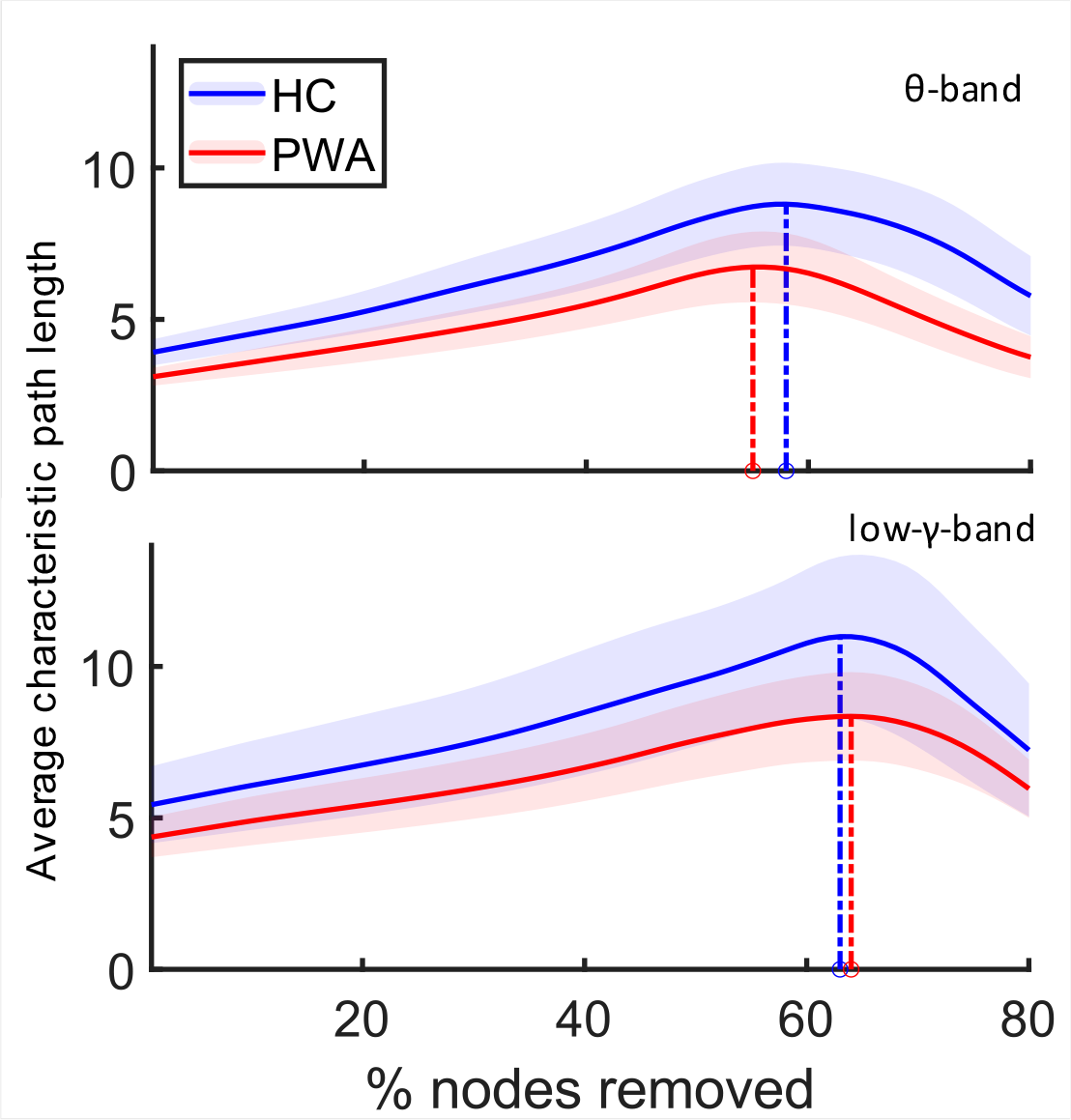
Outcome of the targeted node-attack analysis. For the theta (θ)-band-network, the average characteristic path length showed an earlier peak in PWA compared with HC, whilst no difference emerged within the low-gamma (γ)-band network. Shaded areas represent the 95% confidence interval.

### 3.6. Support vector machine

The potential of network measures to discriminate between PWA and HC was assessed by implementing a SVM classifier with leave-one-out cross-validation (Figure 5). The training set included the average connectivity strength of the NBS-detected component (WPLI_NBS_) at each cross-validation step as well as all graph theory metrics (i.e. node strength, clustering coefficient, eccentricity, characteristic path length, small-worldness and modularity) and age. Only the frequency bands which yielded significant between-group differences, i.e. theta-band and low-gamma-band, were tested. The classifier was able to successfully predict the group to which a subject belonged with an accuracy of 78% and area under the receiver operating characteristic (AUROC) curve of 83%. The optimal working point of the classifier was situated at sensitivity 74% and specificity 82%. The most important variable for classification as assessed by the classifier was the WPLI_NBS_ (obtained at *tth* = 12) in the theta-band, followed by the WPLI_NBS_ (obtained at *tth* = 8) in the low-gamma-band, the node strength in the theta-band, and the characteristic path length in the low-gamma-band. The complete importance ranking with respective coefficients is reported in Supplementary materials.

**Figure 5.**
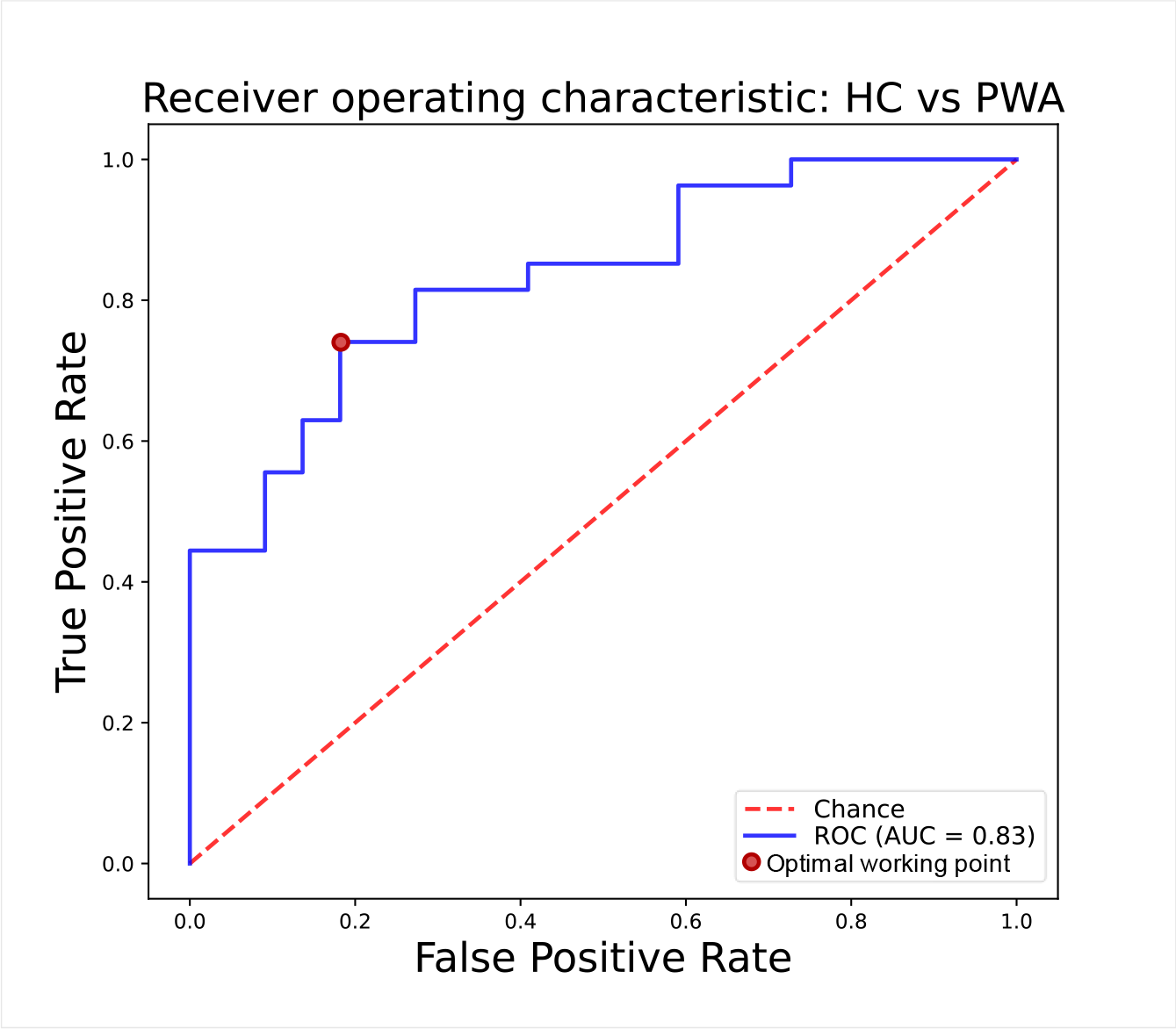
Outcome of the SVM classifier. The leave-one-out cross-validation resulted in AUC = 83%, with optimal working point at sensitivity = 74% and specificity = 82%.

## 4. Discussion

We used EEG to investigate functional network correlates of natural speech processing in PWA. Through a combination of complementary analyses, we provided a picture of the altered synchronization between scalp regions due to the pathological condition. Using NBS, we detected increased connectivity in aphasia between the left temporal area and the right hemisphere in the theta-band, as well as a weakened posterior-frontal subnetwork in the low-gamma-band. The connectivity strength was also negatively correlated with the semantic fluency in PWA in both delta-and low-gamma-band networks. Connectivity changes were reflected into the network geometry as demonstrated with graph theory analysis. In fact, a global reorganization of the network architecture in PWA emerged when compared to HC in the theta-band, whilst only local changes were found in the low-gamma-band network. Alteration of the network architecture in PWA was further assessed with a targeted node-attack approach, which, together with the lower small-worldness, suggested a tendency towards a more scale-free distribution compared to HC. The suitability of EEG-network measures as biomarkers for chronic aphasia was proved with a SVM classifier, which yielded an accuracy of 78% (AUC = 83%).

### 4.1. Between-region synchronization is higher in the theta-band network and lower in the low-gamma-band network in PWA compared to HC

The between-group comparison revealed that aphasia is associated with altered EEG network properties within the theta-band and low-gamma-band. From our analysis, an increased synchronization between the right hemisphere and the left temporal region emerges within the theta-band network in aphasia. In contrast, PWA showed a weakened posterior-anterior network pattern in the low-gamma-band compared to HC.

The role of these two frequency bands can be interpreted in the perspective of the Asymmetric Sampling in Time (AST) theory (Poeppel, 2003). According to the proposed model, the left auditory cortex is involved in the decoding of phonemes, whilst slower acoustic modulations such as speech prosody (Belin et al., 2004) are usually integrated in the right auditory cortex. Processing of phonemes and prosody were shown to be respectively associated with gamma-band and theta-band tracking during natural speech listening (di Liberto et al., 2015; Ding & Simon, 2014; Giraud et al., 2007). Our results suggest that these mechanisms might be functionally affected in PWA.

The existing literature and the AST theory nicely fit with the outcome of our analysis in the theta-band network. As discussed in a review by Hartwigsen & Saur (2019), over-activation of the right hemisphere in domain-general areas at rest or while performing a language test in chronic aphasia is thought to be associated with either a compensatory mechanism (Abel et al., 2015; Breier et al., 2007; Sims et al., 2016) or maladaptive inhibition of the still functioning perilesional regions, as also observed in transcranial magnetic stimulation studies with both healthy participants and persons with acute or chronic aphasia (Dafotakis et al., 2008; Khedr et al., 2014; Naeser et al., 2004). Our approach allows to further disentangle this aspect and favors the former speculation while extending it to natural speech processing. In fact, we detected hyper-synchronization between left and right hemispheric regions rather than a stand-alone over-engagement of the contralesional hemisphere. We believe that this may reflect a recruitment of intact domain-general and homologous regions over the right hemisphere by the damaged left-hemispheric language and auditory areas. In line with our finding, an increase of theta-band activity in both acute and chronic aphasia was observed in a region-of-interest- (ROI-) based EEG study (Kawano et al., 2021), emerging as a negative shift in the power-spectrum of the alpha-band activity (Dalton et al., 2021; Finnigan & van Putten, 2013; P. P. Shah-Basak et al., 2020).

On the other hand, a biological interpretation of gamma-band activity remains challenging (Buzsáki & Wang, 2012; Yuval-Greenberg et al., 2008). High-frequency oscillations and coupled activity with the theta-band are reportedly reflecting networks of hippocampal inhibitory interneurons (Bartos et al., 2007; Herrmann & Demiralp, 2005) associated with memory, attentive and learning processes (Bartos et al., 2007; Belluscio et al., 2012; Colgin et al., 2009; Kay, 2003). Interestingly, these cognitive functions were recently shown to be intact in PWA (Schevenels et al., 2022), leading to the speculation that protective or compensatory mechanisms might be reflected in weakened low-gamma-band network. Our results may seem in contrast with previous studies reporting either stronger connectivity (Snyder et al., 2021) or alteration in alpha- and beta-band networks (P. Shah-Basak et al., 2022) in chronic aphasia. This inconsistency is likely to be due to the chosen measure of connectivity strength, i.e., WPLI vs amplitude or power envelope correlation, and possibly the experimental paradigm, i.e. natural speech listening vs resting state. Therefore, we believe that our results provide further insights into the physiological processes associated with language processing in aphasia rather than contrasting evidence with respect to the existing literature.

### 4.2. EEG connectivity strength correlates with the semantic fluency in aphasia

By means of NBS, we found two network components in PWA negatively correlated with the semantic fluency score, respectively in the delta-band and the low-gamma-band. In the delta-band, the detected component comprised a cluster over the left temporo-parietal region, likely involving the language area, as well as frontal inter-hemispheric connectivity. Previous research reported a correlation between decrease of cortical activity and speech fluency, whilst there is still contrasting evidence from the network perspective (Fridriksson et al., 2012, 2015; P. Shah-Basak et al., 2022; Woodhead et al., 2017). A recent EEG study demonstrated an association between the delta-band activity and the perception of rhythmic properties of speech (Boucher et al., 2019). Hence, a possible interpretation of our result is that PWA with partial recovery retain an altered processing of articulated sounds, which reflects into the semantic fluency; delta-band activity might be dependent on the level of such impairment and would emerge as a network cluster over the involved cortical regions as observed here. However, the reason why worse performance is associated with stronger connectivity remains speculative. As per the between-group analysis, we propose an enhanced cortical recruitment in the perilesional regions aimed at a compensatory mechanism. On the other hand, the correlation between low-gamma-band connectivity over a posterior-anterior pattern and semantic fluency poses a challenge in its interpretation. As discussed above, highest frequency activity is reportedly associated with phonemic processing, however it is unclear how phonemic processing would relate to semantic fluency, and further studies will be needed to investigate this aspect.

An additional concern lies in the fact that semantic fluency is in part related to executive function rather than language processing (Aita et al., 2018). Therefore, we may have not observed network alterations associated with language-related impairment, rather a potential cognitive dysfunction due to the stroke. Recent studies aimed at assessing the behavioral correlates of affected semantic fluency (Chami et al., 2021; Lonie et al., 2009; Whiteside et al., 2015); the reported results, together with the topographical output of NBS (Figure 2), suggest that the correlation between EEG-network metrics and semantic fluency is likely reflecting aphasia-related language processing alterations.

Interestingly, similar findings in chronic aphasia were not reported when amplitude correlation was used as connectivity metric (P. Shah-Basak et al., 2022). In fact, a positive correlation between the alpha- and beta-band network connectivity with fluency score was found, whilst no significant results emerged in other frequency bands. As discussed by the authors, this poses an issue on generalization of results, since analysis outcomes may depend on the connectivity metrics (Colclough et al., 2016).

Surprisingly, no correlation emerged with other behavioral measures, i.e. phonological fluency, naming performance or the ScreeLing test score. As we speculated in the paragraph above, failure to capture functional features associated with these other altered behavioral measures might also be due to a reduced sensitivity of WPLI as connectivity metric. This aspect remains unsolved and should be investigated in future research.

### 4.3. Network geometrical properties are altered globally in the theta-band but only locally in the low-gamma-band

By means of graph theory analysis, we detected an overall rearrangement of the functional network architecture in individuals with chronic aphasia.

Within the theta-band, the patterns of alteration of the node strength in PWA, both globally and locally, concur with the altered connectivity patterns detected with the NBS. Most connections are re- routed in PWA and link most areas with the left language and auditory regions, resulting in a functional centralization of the network towards the lesioned areas, possibly for compensatory purposes. This interpretation is further supported by the increased integration of the network as reflected in the reduced characteristic path length and eccentricity, and by the reduced randomness of the node distribution of the network nodes as measured with lower small-worldness. This latter property already emerges at the acute stage, as shown in a previous graph theory study in resting state where authors reported reduced small-worldness in the theta-band network only in patients with a left-hemispheric lesion (Caliandro et al., 2017). Together with locally increased node strength, we believe this to reflect a redistribution of the network towards a more scale-free distribution while processing natural speech, as also explored with the targeted node-attack approach discussed below. Despite this rearrangement, the modular distribution is not significantly altered in PWA compared to HC in either frequency bands. A recent longitudinal fMRI study investigated the variations of modularity across recovery; at the acute stage, the brain is less modular in aphasia compared to the healthy condition, whilst the functional network tends to a more normative modular distribution towards the chronic stage (Siegel et al., 2018). Our results concur with this latter mechanism, and further prove the efficiency of EEG as a method for inferring functional brain network properties in health and disease.

In contrast, no consistent global alteration of graph metrics was detected in the low-gamma-band in PWA, although local changes emerged which spatially resembled the findings in the theta-band network. As for the theta-band network changes, we propose that local variations in the low-gamma-band network architecture should be interpreted in the perspective of compensatory mechanisms by recruitment of residual intact areas. In fact, higher node strength and clustering coefficient in PWA as obtained with WPLI reflects localized stronger synchronization with other brain regions, i.e. coordinated cortical response to the experimental task. The lack of globally distributed changes within the gamma-band network should not be surprising. In fact, whilst theta-band oscillations reportedly have an integration role across long-range-distributed neuronal populations, gamma-band activity mostly reflects local intraregional neuronal activity, and a coupling between the two can be observed in specific processes, as described in section 4.1 (Lopes da Silva, 2013; Mizuhara et al., 2004; von Stein & Sarnthein, 2000).

### 4.4. A scale-free theta-band network distribution is more prominent in PWA compared to HC

By means of an iterative node-attack, we found a tendency towards a negative shift of the trend of the characteristic path length in PWA with respect to the percentage of removed nodes. In previous studies, the characteristic path length showed an increasing trend, followed by a decrease after reaching a maximum (Barabasi & Albert, 1999; Kaiser et al., 2007; Mehraram et al., 2020; Stam et al., 2009). Reportedly, the maximum peak occurs earlier in scale-free networks when compared with small-world or random networks (Kaiser, et al., 2007). Scale-free networks are less robust to targeted external attacks, due to their higher centralization over a smaller number of nodes, whose distribution follows a power law. There is still a debate on whether the human brain shows a scale-free architecture (Eguiluz et al., 2005; Kaiser et al., 2007). In fact, due to their localized centralization, scale-free networks are generally more robust to random damage, such as lesions or atrophy, than small-world or random networks. Although a healthy brain has been shown to have scale-free properties (Eguiluz et al., 2005), our results suggest aphasia to be associated with a more emphasized scale-free architecture when compared to controls. Together with higher localized node strength and reduced small-worldness, the outcome of the targeted node-attack further suggests that with aphasia the brain tends to centralize the network towards the affected region in order to process natural speech, perhaps for it to recruit other functioning areas as a compensatory mechanism rather than as a maladaptive process.

### 4.5. Connectivity strength is the most discriminant variable between PWA and HC

To test the diagnostic potential of EEG network metrics, we implemented a SVM classifier. Network metrics which significantly differed between groups were included as features. Age was also included to take in account of age-related network alterations (Damoiseaux, 2017; Geerligs et al., 2015), which might reflect into the accuracy of the classifier. Interestingly, despite our limited sample size, our EEG- network-based classifier resulted in an accuracy score of 78%, that is higher than what was obtained with a lesion-based classification of aphasia (Landrigan et al., 2021), and slightly lower than an EEG-power-based classifier for general functional outcome of ischemic stroke (Bentes et al., 2018). Notably, the most discriminant variable was the connectivity strength in both frequency bands. This outcome agrees with previous research on other pathological conditions (Mehraram et al., 2020; Peraza et al., 2018; Xu et al., 2016), and due to its immediate computation compared to other network metrics, this result supports EEG connectivity as a suitable biomarker for aphasia. Further research will be needed to investigate whether EEG network metrics remain efficient to predict the progress of the condition when measured at the (sub)acute stage.

### 4.6. Limitations

Despite its practicality and diagnostic suitability, the main limitation of our study lies in the use of EEG metrics. In fact, electrical signals recorded from the scalp remain challenging to interpret, as they correspond to the spatial linear combination of cortical activations which are normally affected by conducting properties of the surrounding tissues (Michel & He, 2019). This even more so holds in the case of a stroke, where an effect of the lesion properties on the cortical activity should be expected (Cassidy et al., 2020; Park et al., 2016). Nevertheless, we argue that potential conductivity alteration due to the lesion did not alter the significance of the obtained results. Lesion-related alterations of EEG activity would be significant at the group level in the unlikely case that all PWA exhibited the same stroke characteristics. In contrast, given the high variability aphasia (Lazar & Antoniello, 2008), conduction alteration introduces a confounding factor into the statistics. This may potentially hamper the emergence of significant differential network components between groups. Nevertheless, by using the NBS we detected a consistent altered network across PWA, hence we are confident that our statistical approach allowed us to capture the backbone functional networks associated with chronic aphasia. The effect of the lesion on the EEG metrics may be investigated by including an additional participant cohort of stroke patients without aphasia. However, this would pose a challenge in controlling for the variability of lesion characteristics across the cohort. A more effective strategy consists of source-level analysis, where an accurate head model is crucial in order to correctly infer the cortical activity and avoid lesion-related distortions in the measured signal (Piastra et al., 2021, 2022).

Aphasia is often associated with cognitive impairment, as also emerged from the demographic scores presented in the present study. Hence, the possibility exists that part of our findings may be due to differences in cognitive functions between groups rather than solely language impairment. A common approach in clinical studies consists of including a cognitive score as a nuisance covariate. However, according to statistical theory this is not the recommended strategy to this scope (Field et al., 2012). Alternatively, a matched dataset of PWA without clinical impairment might be included at the recruitment stage. However, such approach is not feasible with the available cohort due to the intrinsic characteristics of aphasia, which involve a level of alteration of cognitive functions. Nevertheless, we aimed at reducing the effect of stroke-related cognitive impairment in the statistical analysis by implementing an experimental paradigm based on natural speech listening.

One participants from the PWA group fell asleep for short periods of time during the first three blocks of the story. Given the shortness of the amount of affected recording, we decided to preserve the full EEG of the participant for our analysis. An effect of sleep on functional network properties was reported in the literature, and might be expected to have affected our analysis. Namely, sleep is reportedly associated with a reorganization of the network towards a small-world topology (Ferri et al., 2008; Vecchio et al., 2017) and stronger connectivity in delta and alpha bands (Huang et al., 2021). Nonetheless, we report lower small-worldness in PWA compared to HC and no between-group differences in alpha or delta bands, further proving the robustness of our results and the lack of any bias.

From the methodological perspective, by definition WPLI does not provide any directionality information on the measured connections. Future work should use alternative metrics to infer whether aphasia is associated with direction-specific connectivity changes. Moreover, to control for conducting signals it rejects all zero-lag connections, potentially omitting any true zero-lag connectivity between scalp-regions. Therefore, we believe that our results should be interpreted in comparison and complementarity to other studies which opted for different connectivity measures, such as amplitude correlation (P. Shah-Basak et al., 2022) or Granger causality (Sarmukadam & Behroozmand, 2022).

A limitation of the NBS approach lies in the choice of the primary statistical threshold, which remains arbitrary (Zalesky et al., 2010). As a workaround, we chose several values for each test and reported the outcomes of the analysis for the tested ranges.

Due to the number of statistical tests and inter-dependence of EEG network metrics, correction for multiple comparisons remains challenging. In fact, most methods in the literature involve conservative assumptions which networks and more generally EEG measures fail to meet due to their intrinsic properties. Nevertheless, we aimed at an exploratory analysis to provide an overall picture of disease-related alterations in a comprehensive number of metrics, rather than investigate a specific hypothesis.

## 5. Conclusion

We performed an exploratory investigation of altered EEG functional network metrics associated with aphasia and natural speech perception. We found an overall rearrangement of the functional network, with an enhanced centralization over the lesioned language region, possibly due to compensatory mechanisms associated with poorer recovery outcomes. Our interpretation is further supported by the outcome of the correlation analysis between clinical scores and functional connectivity, which was stronger for poorer performance. Among all network metrics, connectivity strength contributed the most to an effective discrimination between PWA and HC, proving EEG-network connectivity as a suitable biomarker for aphasia at the chronic stage. Further research is needed to assess the cortical sources of the detected abnormalities in the scalp-recorded signal. Furthermore, the potential of EEG network metrics obtained at the acute stage to predict the disease outcomes at the chronic stage remains of clinical interest and should be investigated.

## References

1. Abbas, A. K., Azemi, G., Amiri, S., Ravanshadi, S., & Omidvarnia, A. (2021). Effective connectivity in brain networks estimated using EEG signals is altered in children with ADHD. Computers in Biology and Medicine, 134, 104515. https://doi.org/10.1016/j.compbiomed.2021.104515

2. Abel, S., Weiller, C., Huber, W., Willmes, K., & Specht, K. (2015). Therapy-induced brain reorganization patterns in aphasia. Brain, 138(4), 1097–1112. https://doi.org/10.1093/BRAIN/AWV022

3. Acharya, U. R., Vinitha Sree, S., Swapna, G., Martis, R. J., & Suri, J. S. (2013). Automated EEG analysis of epilepsy: A review. Knowledge-Based Systems, 45, 147–165. https://doi.org/10.1016/J.KNOSYS.2013.02.014

4. Aita, S. L., Beach, J. D., Taylor, S. E., Borgogna, N. C., Harrell, M. N., & Hill, B. D. (2018). Executive, language, or both? An examination of the construct validity of verbal fluency measures. 26(5), 441–451. https://doi.org/10.1080/23279095.2018.1439830

5. Barabasi, A. L., & Albert, R. (1999). Emergence of scaling in random networks. Science, 286(5439), 509–512.

6. Bartos, M., Vida, I., & Jonas, P. (2007). Synaptic mechanisms of synchronized gamma oscillations in inhibitory interneuron networks. Nature Reviews Neuroscience, 8(1), 45–56. https://doi.org/10.1038/nrn2044

7. Behroozmand, R., Bonilha, L., Rorden, C., Hickok, G., & Fridriksson, J. (2022). Neural correlates of impaired vocal feedback control in post-stroke aphasia. NeuroImage, 250(September 2021), 118938. https://doi.org/10.1016/j.neuroimage.2022.118938

8. Belin, P., Fecteau, S., & Bédard, C. (2004). Thinking the voice: neural correlates of voice perception. Trends in Cognitive Sciences, 8(3), 129–135. https://doi.org/10.1016/J.TICS.2004.01.008

9. Belluscio, M. A., Mizuseki, K., Schmidt, R., Kempter, R., & Buzsáki, G. (2012). Cross-Frequency Phase– Phase Coupling between Theta and Gamma Oscillations in the Hippocampus. The Journal of Neuroscience, 32(2), 423–435. https://doi.org/10.1523/jneurosci.4122-11.2012

10. Bentes, C., Peralta, A. R., Viana, P., Martins, H., Morgado, C., Casimiro, C., Franco, A. C., Fonseca, A. C., Geraldes, R., Canhão, P., Pinho e Melo, T., Paiva, T., & Ferro, J. M. (2018). Quantitative EEG and functional outcome following acute ischemic stroke. Clinical Neurophysiology, 129(8), 1680–1687. https://doi.org/10.1016/j.clinph.2018.05.021

11. Bhattacharya, B. sen, Coyle, D., & Maguire, L. P. (2011). A thalamo–cortico–thalamic neural mass model to study alpha rhythms in Alzheimer’s disease. Neural Networks, 24(6), 631–645. https://doi.org/10.1016/j.neunet.2011.02.009

12. Bohr, I. J., Kenny, E., Blamire, A., O’Brien, J. T., Thomas, A. J., Richardson, J., & Kaiser, M. (2013). Resting-state functional connectivity in late-life depression: higher global connectivity and more long distance connections. Frontiers in Psychiatry, 3, 116. https://doi.org/10.3389/fpsyt.2012.00116

13. Boucher, V. J., Gilbert, A. C., & Jemel, B. (2019). The Role of Low-frequency Neural Oscillations in Speech Processing: Revisiting Delta Entrainment. https://doi.org/10.1162/jocn_a_01410

14. Breier, J. I., Maher, L. M., Schmadeke, S., Hasan, K. M., & Papanicolaou, A. C. (2007). Changes in Language-specific Brain Activation after Therapy for Aphasia using Magnetoencephalography: A Case Study. http://Dx.Doi.Org/10.1080/13554790701448200, 13(3), 169–177. https://doi.org/10.1080/13554790701448200

15. Brodbeck, C., Presacco, A., & Simon, J. Z. (2018). Neural source dynamics of brain responses to continuous stimuli: Speech processing from acoustics to comprehension. NeuroImage, 172(December 2017), 162–174. https://doi.org/10.1016/j.neuroimage.2018.01.042

16. Brodbeck, C., & Simon, J. Z. (2020). Continuous speech processing. Current Opinion in Physiology, 18, 25–31. https://doi.org/10.1016/j.cophys.2020.07.014

17. Brownsett, S. L. E., Warren, J. E., Geranmayeh, F., Woodhead, Z., Leech, R., & Wise, R. J. S. (2014). Cognitive control and its impact on recovery from aphasic stroke. Brain, 137(1), 242–254. https://doi.org/10.1093/brain/awt289

18. Buzsáki, G., & Wang, X.-J. (2012). Mechanisms of gamma oscillations. Annual Review of Neuroscience, 35, 203–225.

19. Caliandro, P., Vecchio, F., Miraglia, F., Reale, G., della Marca, G., la Torre, G., Lacidogna, G., Iacovelli, C., Padua, L., Bramanti, P., & Rossini, P. M. (2017). Small-World Characteristics of Cortical Connectivity Changes in Acute Stroke. Neurorehabilitation and Neural Repair, 31(1), 81–94. https://doi.org/10.1177/1545968316662525

20. Canuet, L., Ishii, R., Pascual-Marqui, R. D., Iwase, M., Kurimoto, R., Aoki, Y., Ikeda, S., Takahashi, H., Nakahachi, T., & Takeda, M. (2011). Resting-state EEG source localization and functional connectivity in schizophrenia-like psychosis of epilepsy. PLOS ONE, 6(11), e27863. https://doi.org/10.1371/journal.pone.0027863

21. Cassidy, J. M., Wodeyar, A., Wu, J., Kaur, K., Masuda, A. K., Srinivasan, R., & Cramer, S. C. (2020). Low-Frequency Oscillations Are a Biomarker of Injury and Recovery after Stroke. Stroke, 1442–1450. https://doi.org/10.1161/STROKEAHA.120.028932

22. Chami, S., Charalambous, C., Knijnik, S. R., & Docking, K. (2021). Language and executive function skills as predictors of semantic fluency performance in pre-school children. https://doi.org/10.1080/17549507.2021.2008005.

23. Chen, X., Chen, L., Zheng, S., Wang, H., Dai, Y., Chen, Z., & Huang, R. (2021). Disrupted Brain Connectivity Networks in Aphasia Revealed by Resting-State fMRI. Frontiers in Aging Neuroscience, 13(October), 1–10. https://doi.org/10.3389/fnagi.2021.666301

24. Colclough, G. L., Woolrich, M. W., Tewarie, P. K., Brookes, M. J., Quinn, A. J., & Smith, S. M. (2016). How reliable are MEG resting-state connectivity metrics? Neuroimage, 138, 284–293.

25. Colgin, L. L., Denninger, T., Fyhn, M., Hafting, T., Bonnevie, T., Jensen, O., Moser, M.-B., & Moser, E. I. (2009). Frequency of gamma oscillations routes flow of information in the hippocampus. Nature, 462(7271), 353–357. https://doi.org/10.1038/nature08573

26. Dafotakis, M., Grefkes, C., Wang, L., Fink, G. R., & Nowak, D. A. (2008). The effects of 1 Hz rTMS over the hand area of M1 on movement kinematics of the ipsilateral hand. Journal of Neural Transmission, 115(9), 1269–1274. https://doi.org/10.1007/S00702-008-0064-1/FIGURES/2

27. Dalton, S. G. H., Cavanagh, J. F., & Richardson, J. D. (2021). Spectral Resting-State EEG (rsEEG) in Chronic Aphasia Is Reliable, Sensitive, and Correlates With Functional Behavior. Frontiers in Human Neuroscience, 15(March), 1–16. https://doi.org/10.3389/fnhum.2021.624660

28. Damoiseaux, J. S. (2017). Effects of aging on functional and structural brain connectivity. NeuroImage, 160, 32–40. https://doi.org/10.1016/J.NEUROIMAGE.2017.01.077

29. Dauwan, M., van Dellen, E., van Boxtel, L., van Straaten, E. C. W., de Waal, H., Lemstra, A. W., Gouw, A. A., van der Flier, W. M., Scheltens, P., Sommer, I. E., & Stam, C. J. (2016). EEG-directed connectivity from posterior brain regions is decreased in dementia with Lewy bodies: a comparison with Alzheimer’s disease and controls. Neurobiology of Aging, 41(Supplement C), 122–129. https://doi.org/10.1016/j.neurobiolaging.2016.02.017

30. de Cheveigné, A. (2020). ZapLine: A simple and effective method to remove power line artifacts. NeuroImage, 207(October 2019). https://doi.org/10.1016/j.neuroimage.2019.116356

31. Demeyere, N., Riddoch, M. J., Slavkova, E. D., Bickerton, W.-L., & Humphreys, G. W. (2015). The Oxford Cognitive Screen (OCS): validation of a stroke-specific short cognitive screening tool. Psychological Assessment, 27(3), 883–894. https://doi.org/10.1037/PAS0000082

32. di Liberto, G. M., O’Sullivan, J. A., & Lalor, E. C. (2015). Low-frequency cortical entrainment to speech reflects phoneme-level processing. Current Biology, 25(19), 2457–2465. https://doi.org/10.1016/j.cub.2015.08.030

33. Ding, N., & Simon, J. Z. (2014). Cortical entrainment to continuous speech: Functional roles and interpretations. Frontiers in Human Neuroscience, 8(MAY), 1–7. https://doi.org/10.3389/fnhum.2014.00311

34. Eguiluz, V. M., Chialvo, D. R., Cecchi, G. A., Baliki, M., & Apkarian, A. v. (2005). Scale-free brain functional networks. Phys Rev Lett, 94(1), 18102. https://doi.org/10.1103/PhysRevLett.94.018102

35. el Hachioui, H., van de Sandt-Koenderman, M. W. M. E., Dippel, D. W. J., Koudstaal, P. J., & Visch-Brink, E. G. (2012). The ScreeLing: occurrence of linguistic deficits in acute aphasia post-stroke. Journal of Rehabilitation Medicine, 44(5), 429–435. https://doi.org/10.2340/16501977-0955

36. Ferri, R., Rundo, F., Bruni, O., Terzano, M. G., & Stam, C. J. (2008). The functional connectivity of different EEG bands moves towards small-world network organization during sleep. Clinical Neurophysiology, 119(9), 2026–2036. https://doi.org/10.1016/J.CLINPH.2008.04.294

37. Field, A., Miles, J., & Field, Z. (2012). Discovering statistics using R. Sage London.

38. Finnigan, S., & van Putten, M. J. A. M. (2013). EEG in ischaemic stroke: Quantitative EEG can uniquely inform (sub-)acute prognoses and clinical management. Clinical Neurophysiology, 124(1), 10– 19. https://doi.org/10.1016/J.CLINPH.2012.07.003

39. Flamand-Roze, C., Falissard, B., Roze, E., Maintigneux, L., Beziz, J., Chacon, A., Join-Lambert, C., Adams, D., & Denier, C. (2011). Validation of a new language screening tool for patients with acute stroke: the Language Screening Test (LAST). Stroke, 42(5), 1224–1229.

40. Florin, E., Gross, J., Pfeifer, J., Fink, G. R., & Timmermann, L. (2010). The effect of filtering on Granger causality based multivariate causality measures. NeuroImage, 50(2), 577–588. https://doi.org/10.1016/j.neuroimage.2009.12.050

41. Flowers, H. L., Skoretz, S. A., Silver, F. L., Rochon, E., Fang, J., Flamand-Roze, C., & Martino, R. (2016). Poststroke Aphasia Frequency, Recovery, and Outcomes: A Systematic Review and Meta-Analysis. Archives of Physical Medicine and Rehabilitation, 97(12), 2188–2201.e8. https://doi.org/10.1016/j.apmr.2016.03.006

42. Fluss, R., Faraggi, D., & Reiser, B. (2005). Estimation of the Youden Index and its associated cutoff point. Biometrical Journal: Journal of Mathematical Methods in Biosciences, 47(4), 458–472.

43. Francart, T., van Wieringen, A., & Wouters, J. (2008). APEX 3: a multi-purpose test platform for auditory psychophysical experiments. Journal of Neuroscience Methods, 172(2), 283–293. https://doi.org/10.1016/j.jneumeth.2008.04.020

44. Fridriksson, J., Basilakos, A., Hickok, G., Bonilha, L., & Rorden, C. (2015). Speech entrainment compensates for Broca’s area damage. Cortex, 69, 68–75. https://doi.org/10.1016/J.CORTEX.2015.04.013

45. Fridriksson, J., Hubbard, H. I., Hudspeth, S. G., Holland, A. L., Bonilha, L., Fromm, D., & Rorden, C. (2012). Speech entrainment enables patients with Broca’s aphasia to produce fluent speech. Brain, 135(12), 3815–3829. https://doi.org/10.1093/BRAIN/AWS301

46. Geerligs, L., Renken, R. J., Saliasi, E., Maurits, N. M., & Lorist, M. M. (2015). A Brain-Wide Study of Age-Related Changes in Functional Connectivity. Cerebral Cortex, 25(7), 1987–1999. https://doi.org/10.1093/CERCOR/BHU012

47. Gialanella, B., Bertolinelli, M., Lissi, M., & Prometti, P. (2010). Predicting outcome after stroke: the role of aphasia. 33(2), 122–129. https://doi.org/10.3109/09638288.2010.488712

48. Giraud, A. L., Kleinschmidt, A., Poeppel, D., Lund, T. E., Frackowiak, R. S. J., & Laufs, H. (2007). Endogenous Cortical Rhythms Determine Cerebral Specialization for Speech Perception and Production. Neuron, 56(6), 1127–1134. https://doi.org/10.1016/j.neuron.2007.09.038

49. Hagoort, P., Brown, C. M., & Swaab, T. Y. (1996). Lexical—semantic event–related potential effects in patients with left hemisphere lesions and aphasia, and patients with right hemisphere lesions without aphasia. Brain, 119(2), 627–649. https://doi.org/10.1093/BRAIN/119.2.627

50. Hamilton, L. S., & Huth, A. G. (2018). The revolution will not be controlled: natural stimuli in speech neuroscience. *Language*, Cognition and Neuroscience, 35(5), 573–582. https://doi.org/10.1080/23273798.2018.1499946

51. Hartwigsen, G., & Saur, D. (2019). Neuroimaging of stroke recovery from aphasia – Insights into plasticity of the human language network. NeuroImage, 190(August 2017), 14–31. https://doi.org/10.1016/j.neuroimage.2017.11.056

52. Herrmann, C. S., & Demiralp, T. (2005). Human EEG gamma oscillations in neuropsychiatric disorders.Clinical Neurophysiology, 116(12), 2719–2733. https://doi.org/10.1016/j.clinph.2005.07.007

53. Hill, A. T., Zomorrodi, R., Hadas, I., Farzan, F., Voineskos, D., Throop, A., Fitzgerald, P. B., Blumberger, D. M., & Daskalakis, Z. J. (2021). Resting-state electroencephalographic functional network alterations in major depressive disorder following magnetic seizure therapy. Progress in Neuro- Psychopharmacology and Biological Psychiatry, 108, 110082. https://doi.org/10.1016/j.pnpbp.2020.110082

54. Huang, H., Zhang, J., Zhu, L., Tang, J., Lin, G., Kong, W., Lei, X., & Zhu, L. (2021). EEG-Based Sleep Staging Analysis with Functional Connectivity. Sensors 2021, *Vol. 21, Page* 1988, *21*(6), 1988. https://doi.org/10.3390/S21061988

55. Kaiser, M. (2011). A tutorial in connectome analysis: Topological and spatial features of brain networks. Neuroimage, 57(3), 892–907. https://doi.org/10.1016/j.neuroimage.2011.05.025

56. Kaiser, M., Martin, R., Andras, P., & Young, M. P. (2007). Simulation of robustness against lesions of cortical networks. European Journal of Neuroscience, 25(10), 3185–3192. https://doi.org/doi:10.1111/j.1460-9568.2007.05574.x

57. Kawano, T., Hattori, N., Uno, Y., Hatakenaka, M., Yagura, H., Fujimoto, H., Nagasako, M., Mochizuki, H., Kitajo, K., & Miyai, I. (2021). Association between aphasia severity and brain network alterations after stroke assessed using the electroencephalographic phase synchrony index. Scientific Reports, 11(1), 1–14. https://doi.org/10.1038/s41598-021-91978-7

58. Kay, L. M. (2003). Two species of gamma oscillations in the olfactory bulb: dependence on behavioral state and synaptic interactions. Journal of Integrative Neuroscience, 2(1), 31–44. https://doi.org/10.1142/S0219635203000196

59. Khedr, E. M., Abo El-Fetoh, N., Ali, A. M., El-Hammady, D. H., Khalifa, H., Atta, H., & Karim, A. A. (2014). Dual-hemisphere repetitive transcranial magnetic stimulation for rehabilitation of poststroke aphasia: A randomized, double-blind clinical trial. Neurorehabilitation and Neural Repair, 28(8), 740–750. https://doi.org/10.1177/1545968314521009/ASSET/IMAGES/LARGE/10.1177_15459683145210 09-FIG2.JPEG

60. Kiran, S., & Thompson, C. K. (2019). Neuroplasticity of language networks in aphasia: Advances, updates, and future challenges. Frontiers in Neurology, 10(APR). https://doi.org/10.3389/fneur.2019.00295

61. Kong, Y. Y., Somarowthu, A., & Ding, N. (2015). Effects of Spectral Degradation on Attentional Modulation of Cortical Auditory Responses to Continuous Speech. JARO - Journal of the Association for Research in Otolaryngology, 16(6), 783–796. https://doi.org/10.1007/S10162-015-0540-X/FIGURES/7

62. Kries, J., de Clercq, P., Lemmens, R., Francart, T., & Vandermosten, M. (2022). Tuning in on auditory details is difficult: Individuals with aphasia show impaired acoustic and phonemic processing. BioRxiv, 2022.12.14.520503. https://doi.org/10.1101/2022.12.14.520503

63. Landrigan, J. F., Zhang, F., & Mirman, D. (2021). A data-driven approach to post-stroke aphasia classification and lesion-based prediction. Brain, 144(5), 1372–1383. https://doi.org/10.1093/brain/awab010

64. Langer, N., Pedroni, A., & Jancke, L. (2013). The problem of thresholding in small-world network analysis. PLOS ONE, 8(1), e53199. https://doi.org/10.1371/journal.pone.0053199

65. Lazar, R. M., & Antoniello, D. (2008). Variability in recovery from aphasia. Current Neurology and Neuroscience Reports, 8(6), 497–502. https://doi.org/10.1007/S11910-008-0079-X/METRICS

66. Li, R., Mukadam, N., & Kiran, S. (2022). Functional MRI evidence for reorganization of language networks after stroke. Handbook of Clinical Neurology, 185, 131–150. https://doi.org/10.1016/B978-0-12-823384-9.00007-4

67. Lonie, J. A., Herrmann, L. L., Tierney, K. M., Donaghey, C., O’Carroll, R., Lee, A., & Ebmeier, K. P. (2009). Lexical and semantic fluency discrepancy scores in aMCI and early Alzheimer’s disease. Journal of Neuropsychology, 3(1), 79–92. https://doi.org/10.1348/174866408X289935

68. Lopes da Silva, F. (2013). EEG and MEG: relevance to neuroscience. Neuron, 80(5), 1112–1128. https://doi.org/10.1016/j.neuron.2013.10.017

69. Mehraram, R., Kaiser, M., Cromarty, R., Graziadio, S., O’Brien, J. T., Killen, A., Taylor, J. P., & Peraza, L. R. (2020). Weighted network measures reveal differences between dementia types: An EEG study. Human Brain Mapping, 41(6), 1573–1590. https://doi.org/10.1002/hbm.24896

70. Mehraram, R., Peraza, L. R., Murphy, N. R. E., Cromarty, R. A., Graziadio, S., O’Brien, J. T., Killen, A., Colloby, S. J., Firbank, M., Su, L., Collerton, D., Taylor, J.-P., & Kaiser, M. (2022). Functional and structural brain network correlates of visual hallucinations in Lewy body dementia. Brain, 2190–2205. https://doi.org/10.1093/brain/awac094

71. Michel, C. M., & He, B. (2019). EEG source localization. Handbook of Clinical Neurology, 160, 85–101. https://doi.org/10.1016/B978-0-444-64032-1.00006-0

72. Mizuhara, H., Wang, L. Q., Kobayashi, K., & Yamaguchi, Y. (2004). A long-range cortical network emerging with theta oscillation in a mental task. NeuroReport, 15(8), 1233–1238. https://doi.org/10.1097/01.WNR.0000126755.09715.B3

73. Moran, R., Pinotsis, D., & Friston, K. (2013). Neural masses and fields in dynamic causal modeling. Frontiers in Computational Neuroscience, 7. https://www.frontiersin.org/articles/10.3389/fncom.2013.00057

74. Mullen, T. R., Kothe, C. A. E., Chi, Y. M., Ojeda, A., Kerth, T., Makeig, S., Jung, T.-P., & Cauwenberghs, G. (2015). Real-time neuroimaging and cognitive monitoring using wearable dry EEG. IEEE Transactions on Biomedical Engineering, 62(11), 2553–2567.

75. Naeser, M. A., Martin, P. I., Baker, E. H., Hodge, S. M., Sczerzenie, S. E., Nicholas, M., Palumbo, C. L., Goodglass, H., Wingfield, A., Samaraweera, R., Harris, G., Baird, A., Renshaw, P., & Yurgelun-Todd, D. (2004). Overt propositional speech in chronic nonfluent aphasia studied with the dynamic susceptibility contrast fMRI method. NeuroImage, 22(1), 29–41. https://doi.org/10.1016/J.NEUROIMAGE.2003.11.016

76. Nicolo, P., Rizk, S., Magnin, C., Pietro, M. di, Schnider, A., & Guggisberg, A. G. (2015). Coherent neural oscillations predict future motor and language improvement after stroke. Brain, 138(10), 3048– 3060. https://doi.org/10.1093/brain/awv200

77. Onnela, J.-P., Saramäki, J., Kertész, J., & Kaski, K. (2005). Intensity and coherence of motifs in weighted complex networks. Physical Review E, 71(6), 65103.

78. Oostenveld, R., Fries, P., Maris, E., & Schoffelen, J. M. (2011). FieldTrip: Open source software for advanced analysis of MEG, EEG, and invasive electrophysiological data. Comput Intell Neurosci, 2011, 156869. https://doi.org/10.1155/2011/156869

79. Oostenveld, R., & Praamstra, P. (2001). The five percent electrode system for high-resolution EEG and ERP measurements. Clinical Neurophysiology, 112(4), 713–719.

80. Park, W., Kwon, G. H., Kim, Y. H., Lee, J. H., & Kim, L. (2016). EEG response varies with lesion location in patients with chronic stroke. Journal of NeuroEngineering and Rehabilitation, 13(1), 1–10. https://doi.org/10.1186/S12984-016-0120-2/FIGURES/4

81. Pedregosa, F., Varoquaux, G., Gramfort, A., Michel, V., Thirion, B., Grisel, O., Blondel, M., Prettenhofer, P., Weiss, R., Dubourg, V., Vanderplas, J., Passos, A., Cournapeau, D., Brucher, M., Perrot, M., & Duchesnay, É. (2011). Scikit-learn: Machine Learning in Python. The Journal of Machine Learning Research. https://doi.org/10.5555/1953048.2078195

82. Pedroni, A., Bahreini, A., & Langer, N. (2019). Automagic: Standardized preprocessing of big EEG data. NeuroImage, 200(June), 460–473. https://doi.org/10.1016/j.neuroimage.2019.06.046

83. Peraza, L. R., Asghar, A. U. R., Green, G., & Halliday, D. M. (2012). Volume conduction effects in brain network inference from electroencephalographic recordings using phase lag index. J Neurosci Methods, 207(2), 189–199. https://doi.org/10.1016/j.jneumeth.2012.04.007

84. Peraza, L. R., Cromarty, R., Kobeleva, X., Firbank, M. J., Killen, A., Graziadio, S., Thomas, A. J., O’Brien, J. T., & Taylor, J. P. (2018). Electroencephalographic derived network differences in Lewy body dementia compared to Alzheimer’s disease patients. Sci Rep, 8(1), 4637. https://doi.org/10.1038/s41598-018-22984-5

85. Peraza, L. R., Taylor, J.-P., & Kaiser, M. (2015). Divergent brain functional network alterations in dementia with Lewy bodies and Alzheimer’s disease. Neurobiology of Aging, 36(9), 2458–2467. https://doi.org/10.1016/j.neurobiolaging.2015.05.015

86. Perkins, N. J., & Schisterman, E. F. (2005). The Youden index and the optimal cut point corrected for measurement error. Biometrical Journal: Journal of Mathematical Methods in Biosciences, 47(4), 428–441.

87. Piastra, M. C., Oostenveld, R., Schoffelen, J. M., & Piai, V. (2022). Estimating the influence of stroke lesions on MEG source reconstruction. NeuroImage, 260. https://doi.org/10.1016/J.NEUROIMAGE.2022.119422

88. Piastra, M. C., van der Cruijsen, J., Piai, V., Jeukens, F. E. M., Manoochehri, M., Schouten, A. C., Selles, R. W., & Oostendorp, T. (2021). ASH: an Automatic pipeline to generate realistic and individualized chronic Stroke volume conduction Head models. Journal of Neural Engineering, 18(4). https://doi.org/10.1088/1741-2552/ABF00B

89. Pion-Tonachini, L., Kreutz-Delgado, K., & Makeig, S. (2019). ICLabel: An automated electroencephalographic independent component classifier, dataset, and website. NeuroImage, 198, 181–197. https://doi.org/10.1016/j.neuroimage.2019.05.026

90. Poeppel, D. (2003). The analysis of speech in different temporal integration windows: cerebral lateralization as ‘asymmetric sampling in time.’ Speech Communication, 41(1), 245–255. https://doi.org/10.1016/S0167-6393(02)00107-3

91. Pulvermüller, F., Mohr, B., & Lutzenberger, W. (2004). Neurophysiological correlates of word and pseudo-word processing in well-recovered aphasics and patients with right-hemispheric stroke. Psychophysiology, 41(4), 584–591. https://doi.org/10.1111/J.1469-8986.2004.00188.X

92. Rubinov, M., Knock, S. A., Stam, C. J., Micheloyannis, S., Harris, A. W., Williams, L. M., & Breakspear, M. (2009). Small-world properties of nonlinear brain activity in schizophrenia. Hum Brain Mapp, 30(2), 403–416. https://doi.org/10.1002/hbm.20517

93. Rubinov, M., & Sporns, O. (2010). Complex network measures of brain connectivity: Uses and interpretations. Neuroimage, 52(3), 1059–1069. https://doi.org/10.1016/j.neuroimage.2009.10.003

94. Sakkalis, V. (2011). Review of advanced techniques for the estimation of brain connectivity measured with EEG/MEG. Computers in Biology and Medicine, 41(12), 1110–1117. https://doi.org/10.1016/j.compbiomed.2011.06.020

95. Sarmukadam, K., & Behroozmand, R. (2022). Aberrant beta-band brain connectivity predicts speech motor planning deficits in post-stroke aphasia. Cortex, 155, 75–89. https://doi.org/10.1016/J.CORTEX.2022.07.001

96. Saur, D., Lange, R., Baumgaertner, A., Schraknepper, V., Willmes, K., Rijntjes, M., & Weiller, C. (2006). Dynamics of language reorganization after stroke. Brain, 129(6), 1371–1384. https://doi.org/10.1093/brain/awl090

97. Schevenels, K., Michiels, L., Lemmens, R., de Smedt, B., Zink, I., & Vandermosten, M. (2022). The role of the hippocampus in statistical learning and language recovery in persons with post stroke aphasia. NeuroImage: Clinical, 36. https://doi.org/10.1016/J.NICL.2022.103243

98. Schevenels, K., Price, C. J., Zink, I., de Smedt, B., & Vandermosten, M. (2020). A Review on Treatment-Related Brain Changes in Aphasia. Neurobiology of Language, 1(4), 402–433. https://doi.org/10.1162/nol_a_00019

99. Schumacher, J., Thomas, A. J., Peraza, L. R., Firbank, M., Cromarty, R., Hamilton, C. A., Donaghy, P. C., O’Brien, J. T., & Taylor, J.-P. (2020). EEG alpha reactivity and cholinergic system integrity in Lewy body dementia and Alzheimer’s disease. Alzheimer’s Research & Therapy, 12, 1–12.

100. Shah-Basak, P. P., Sivaratnam, G., Teti, S., Francois-Nienaber, A., Yossofzai, M., Armstrong, S., Nayar, S., Jokel, R., & Meltzer, J. (2020). High definition transcranial direct current stimulation modulates abnormal neurophysiological activity in post-stroke aphasia. Scientific Reports, 10(1). https://doi.org/10.1038/S41598-020-76533-0

101. Shah-Basak, P., Sivaratnam, G., Teti, S., Deschamps, T., Kielar, A., Jokel, R., & Meltzer, J. A. (2022). Electrophysiological connectivity markers of preserved language functions in post-stroke aphasia. NeuroImage: Clinical, 34(November 2021), 103036. https://doi.org/10.1016/j.nicl.2022.103036

102. Shin, Y. W., O’Donnell, B. F., Youn, S., & Kwon, J. S. (2011). Gamma Oscillation in Schizophrenia. Psychiatry Investigation, 8(4), 288. https://doi.org/10.4306/PI.2011.8.4.288

103. Siegel, J. S., Ramsey, L. E., Snyder, A. Z., Metcalf, N. v., Chacko, R. v., Weinberger, K., Baldassarre, A., Hacker, C. D., Shulman, G. L., & Corbetta, M. (2016). Disruptions of network connectivity predict impairment in multiple behavioral domains after stroke. Proceedings of the National Academy of Sciences of the United States of America, 113(30), E4367–E4376. https://doi.org/10.1073/pnas.1521083113

104. Siegel, J. S., Seitzman, B. A., Ramsey, L. E., Ortega, M., Gordon, E. M., Dosenbach, N. U. F., Petersen, S. E., Shulman, G. L., & Corbetta, M. (2018). Re-emergence of modular brain networks in stroke recovery. Cortex, 101, 44–59. https://doi.org/10.1016/j.cortex.2017.12.019

105. Sims, J. A., Kapse, K., Glynn, P., Sandberg, C., Tripodis, Y., & Kiran, S. (2016). The relationships between the amount of spared tissue, percent signal change, and accuracy in semantic processing in aphasia. Neuropsychologia, 84, 113–126. https://doi.org/10.1016/j.neuropsychologia.2015.10.019

106. Snyder, D. B., Schmit, B. D., Hyngstrom, A. S., & Beardsley, S. A. (2021). Electroencephalography resting-state networks in people with Stroke. Brain and Behavior, 11(5), 18–35. https://doi.org/10.1002/brb3.2097

107. Soleimani, B., Das, P., Dushyanthi Karunathilake, I. M., Kuchinsky, S. E., Simon, J. Z., & Babadi, B. (2022). NLGC: Network localized Granger causality with application to MEG directional functional connectivity analysis. NeuroImage, 260(June), 119496. https://doi.org/10.1016/j.neuroimage.2022.119496

108. Spironelli, C., & Angrilli, A. (2015). Brain plasticity in aphasic patients: Intra- and inter-hemispheric reorganisation of the whole linguistic network probed by N150 and N350 components. Scientific Reports, 5(October 2014), 1–14. https://doi.org/10.1038/srep12541

109. Stam, C. J., de Haan, W., Daffertshofer, A., Jones, B. F., Manshanden, I., van Cappellen van Walsum, A. M., Montez, T., Verbunt, J. P. A., de Munck, J. C., van Dijk, B. W., Berendse, H. W., & Scheltens, P. (2009). Graph theoretical analysis of magnetoencephalographic functional connectivity in Alzheimer’s disease. Brain, 132(1), 213–224. https://doi.org/10.1093/brain/awn262

110. Taylor, P. N., Papasavvas, C. A., Owen, T. W., Schroeder, G. M., Hutchings, F. E., Chowdhury, F. A., Diehl, B., Duncan, J. S., McEvoy, A. W., Miserocchi, A., de Tisi, J., Vos, S. B., Walker, M. C., & Wang, Y. (2022). Normative brain mapping of interictal intracranial EEG to localize epileptogenic tissue. Brain, 145(3), 939–949. https://doi.org/10.1093/brain/awab380

111. van Ewijk, L., Dijkhuis, L., Hofs-van Kats, M., Hendrickx-Jessurun, M., Wijngaarden, M., & de Hilster, C. (2018). Nederlandse Benoem Test. Springer.

112. Vanthornhout, J., Decruy, L., Wouters, J., Simon, J. Z., & Francart, T. (2018). Speech Intelligibility Predicted from Neural Entrainment of the Speech Envelope. JARO - Journal of the Association for Research in Otolaryngology, 19(2), 181–191. https://doi.org/10.1007/S10162-018-0654-Z/FIGURES/6

113. Vecchio, F., Miraglia, F., Gorgoni, M., Ferrara, M., Iberite, F., Bramanti, P., De Gennaro, L., & Rossini, P. M. (2017). Cortical connectivity modulation during sleep onset: A study via graph theory on EEG data. Human Brain Mapping, 38(11), 5456–5464. https://doi.org/10.1002/HBM.23736

114. Vinck, M., Oostenveld, R., van Wingerden, M., Battaglia, F., & Pennartz, C. M. (2011). An improved index of phase-synchronization for electrophysiological data in the presence of volume-conduction, noise and sample-size bias. Neuroimage, 55(4), 1548–1565. https://doi.org/10.1016/j.neuroimage.2011.01.055

115. Visch-Brink, E., Vandenborre, D., de Smet, H. J., & Mariën, P. (2014). Comprehensive Aphasia Test- Nederlandse bewerking-Handleiding. The Netherlands: Pearson.

116. von Stein, A., & Sarnthein, J. (2000). Different frequencies for different scales of cortical integration: from local gamma to long range alpha/theta synchronization. International Journal of Psychophysiology, 38(3), 301–313. https://doi.org/10.1016/S0167-8760(00)00172-0

117. Whiteside, D. M., Kealey, T., Semla, M., Luu, H., Rice, L., Basso, M. R., & Roper, B. (2015). Verbal Fluency: Language or Executive Function Measure?, 23(1), 29–34. https://doi.org/10.1080/23279095.2015.1004574

118. Woodhead, Z. V. J., Crinion, J., Teki, S., Penny, W., Price, C. J., & Leff, A. P. (2017). Auditory training changes temporal lobe connectivity in ‘Wernicke’s aphasia’: a randomised trial. *Journal of Neurology*, Neurosurgery & Psychiatry, 88(7), 586–594. https://doi.org/10.1136/JNNP-2016-314621

119. Xu, T., Cullen, K. R., Mueller, B., Schreiner, M. W., Lim, K. O., Schulz, S. C., & Parhi, K. K. (2016). Network analysis of functional brain connectivity in borderline personality disorder using resting-state fMRI. Neuroimage Clin, 11, 302–315. https://doi.org/10.1016/j.nicl.2016.02.006

120. Yuval-Greenberg, S., Tomer, O., Keren, A. S., Nelken, I., & Deouell, L. Y. (2008). Transient induced gamma-band response in EEG as a manifestation of miniature saccades. Neuron, 58(3), 429– 441. https://doi.org/10.1016/J.NEURON.2008.03.027

121. Zalesky, A., Fornito, A., & Bullmore, E. T. (2010). Network-based statistic: identifying differences in brain networks. Neuroimage, 53(4), 1197–1207. https://doi.org/10.1016/j.neuroimage.2010.06.041

122. Zhang, G., Si, Y., & Dang, J. (2019). Revealing the Dynamic Brain Connectivity from Perception of Speech Sound to Semantic Processing by EEG. Neuroscience, 415, 70–76. https://doi.org/10.1016/J.NEUROSCIENCE.2019.07.023

123. Zhu, Y., Bai, L., Liang, P., Kang, S., Gao, H., & Yang, H. (2017). Disrupted brain connectivity networks in acute ischemic stroke patients. Brain Imaging and Behavior, 11(2), 444–453. https://doi.org/10.1007/s11682-016-9525-6

