## Supplementary materials for "EEG reveals brain network alterations in chronic aphasia during natural speech listening"

Contents:

1. Graph theory measures
2. Network Based Statistics (NBS): significant outcomes at all threshold values
3. Support vector machine (SVM): variable importance ranking

### 1. Graph theory measures

We investigated the functional network architecture by means of a graph theory study. Metrics of interest included local measures, i.e. node strength, clustering coefficient and eccentricity, and global measures, i.e. average characteristic path length, small-worldness and modularity.

The strength of a node is the sum of the weights of all links connected to the node (Opsahl et al., 2010; Watts & Strogatz, 1998). High connectivity of a region with the rest of the network is reflected into high average node strength of that region.

The clustering coefficient of a node is the ratio between the number of connections and the number of maximum possible connections between neighbors of the node (Kaiser, 2011). We measured the weighted clustering coefficient, which is obtained by weighting the clustering coefficient by the sum of the intensities of the subnetworks formed by the inter-neighbor links (Onnela et al., 2005). The weighted clustering coefficient is sensitive to both number and weights of links between neighbors (Onnela et al., 2005). The (weighted) clustering coefficient spans between 0 (fully disconnected neighbors) and 1 (fully connected neighbors).

Node eccentricity is the longest among the shortest path lengths between the node and all other nodes. Higher network integration reflects into lower average eccentricity, i.e. more short-range connectivity (Utianski et al., 2016).

The characteristic path length is the length of the shortest path between every couple of nodes (Rubinov & Sporns, 2010). In weighted graphs, connection lengths are first computed as inverse weights (Dijkstra, 1959; Newman, 2001), and the Dijkstra algorithm is used to measure all shortest paths (Dijkstra, 1959).

The small-worldness reflects the segregation of the networks as compared to their surrogate random networks (Humphries & Gurney, 2008). For its estimation, the clustering coefficient and the average characteristic path length are normalized by the same measures averaged across 50 surrogate random graphs with the same node degree as the original networks (Humphries & Gurney, 2008; Sanz-Arigita et al., 2010); the small-worldness is obtained as a ratio between the two normalized measures.

The modularity is greater in networks with higher connectivity within communities as compared to connectivity between communities (Newman & Girvan, 2004). Its computation consist of the difference between the fraction of edges within modules and the expected fraction of randomly distributed edges. Network modularity is bounded between -1 and 1.

For a review on network measures and statistics see (Rubinov & Sporns, 2010) and (Kaiser, 2011).

### 2. Network Based Statistics (NBS): significant outcomes at all threshold values

We used the NBS for the between-group network comparison as well as to test correlations between connectivity patterns and clinical test scores. This method requires the arbitrary choice of a primary statistical threshold, depending on how local the effect of interest should be (Zalesky et al., 2010). For both analysis we tested a range of primary thresholds, which we reported in Figure A, Figure B, Figure C and Figure D.

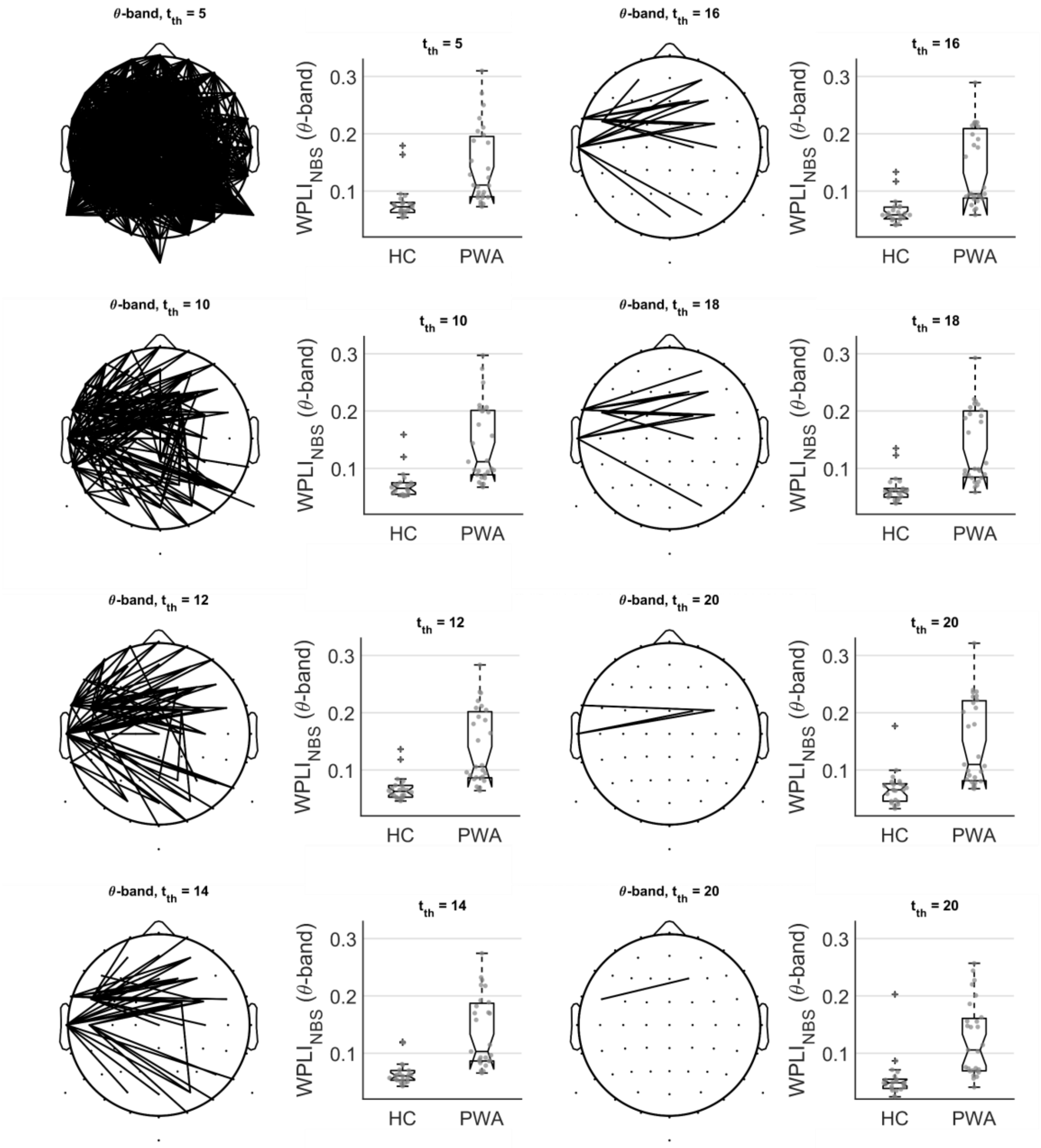


Figure A - NBS: HC vs PWA. Theta-band. Note that for $t_{th}$ = 20, two significant components were detected.

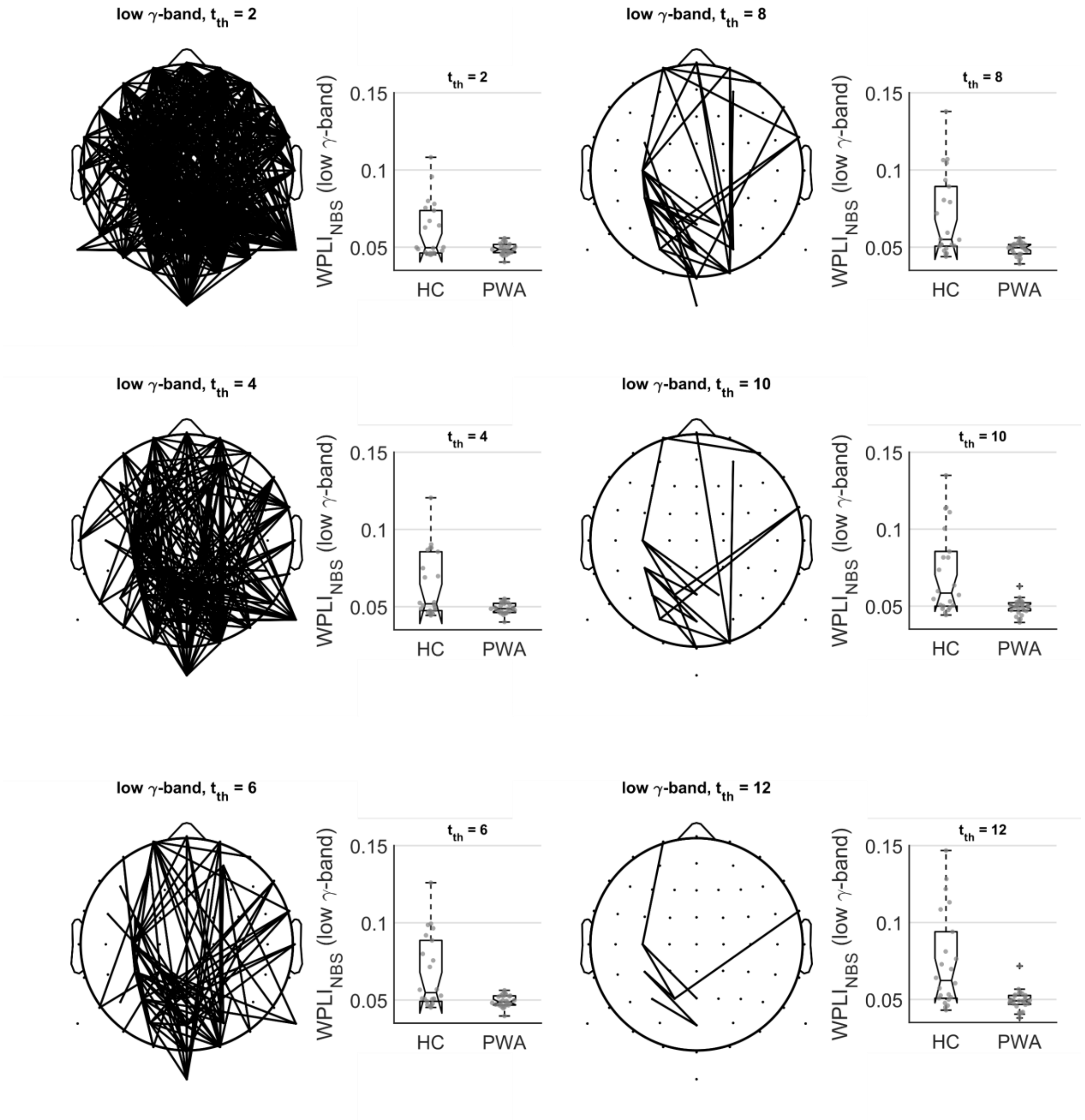


Figure B - NBS: HC vs PWA. Low-gamma-band.

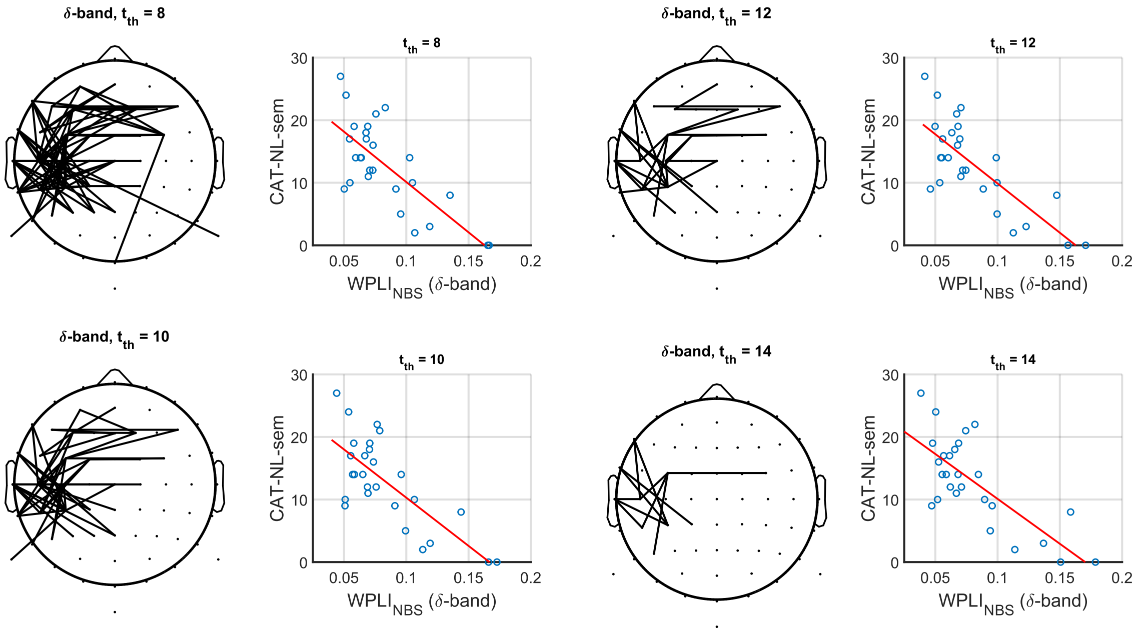


Figure C - NBS: correlation of PWA network with the semantic fluency subscore of the CAT-NL. Delta-band.

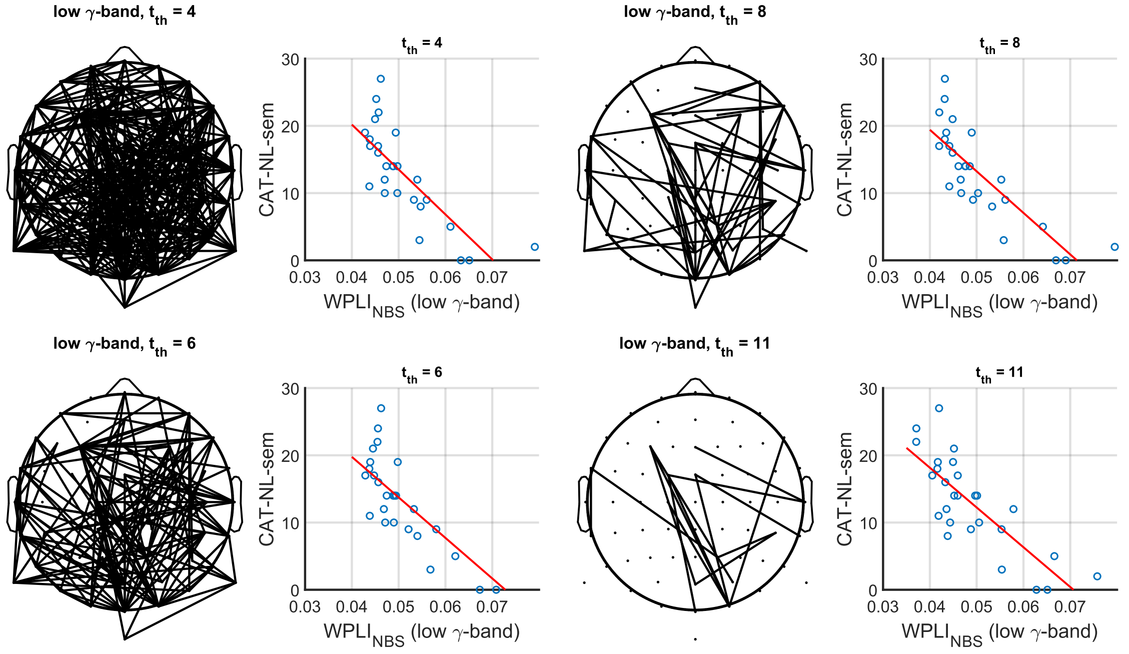


Figure D - NBS: correlation of PWA network with the semantic fluency subscore of the CAT-NL. Low-gamma-band.

### 3. Support vector machine (SVM): variable importance ranking

We implemented an SVM to test the diagnostic potential of EEG network metrics. We found an AUC of 83% to classify the participants as PWA or HC. The variable importance ranking based on their weights is reported in Table A.

Table A - SVM: variable ranking

| Variable | Frequency band | Weight |
|---|---|---|
| $WPLI_{NBS}$ | theta | 2.3182 ± 1.0885 |
| $WPLI_{NBS}$ | low-gamma | 1.9046 ± 1.0776 |
| Node strength | theta | 1.0018 ± 0.6645 |
| Characteristic path length | low-gamma | 0.5815 ± 0.7618 |
| Clustering coefficient | theta | 0.5784 ± 0.0978 |
| Eccentricity | low-gamma | 0.4112 ± 0.2824 |
| Characteristic path length | theta | 0.3044 ± 0.5833 |
| Node strength | low-gamma | 0.2939 ± 0.5353 |
| Eccentricity | theta | 0.1843 ± 0.1393 |
| Clustering coefficient | low-gamma | 0.1012 ± 0.1839 |

#### References


Dijkstra, E. W. (1959). A note on two problems in connexion with graphs. *Numerische Mathematik*, *1*(1), 269–271.

Humphries, M. D., & Gurney, K. (2008). Network 'Small-World-Ness': A Quantitative Method for Determining Canonical Network Equivalence. *PLOS ONE*, *3*(4), e0002051. https://doi.org/10.1371/journal.pone.0002051

Kaiser, M. (2011). A tutorial in connectome analysis: Topological and spatial features of brain networks. *Neuroimage*, *57*(3), 892–907. https://doi.org/https://doi.org/10.1016/j.neuroimage.2011.05.025

Newman, M. E. J. (2001). Scientific collaboration networks. II. Shortest paths, weighted networks, and centrality. *Physical Review E*, *64*(1), 16132.

Newman, M. E. J., & Girvan, M. (2004). Finding and evaluating community structure in networks. *Physical Review E*, *69*(2), 26113. https://doi.org/10.1103/PhysRevE.69.026113

Onnela, J.-P., Saramäki, J., Kertész, J., & Kaski, K. (2005). Intensity and coherence of motifs in weighted complex networks. *Physical Review E*, *71*(6), 65103.

Opsahl, T., Agneessens, F., & Skvoretz, J. (2010). Node centrality in weighted networks: Generalizing degree and shortest paths. *Social Networks*, *32*(3), 245–251. https://doi.org/https://doi.org/10.1016/j.socnet.2010.03.006

Rubinov, M., & Sporns, O. (2010). Complex network measures of brain connectivity: Uses and interpretations. *Neuroimage*, *52*(3), 1059–1069. https://doi.org/https://doi.org/10.1016/j.neuroimage.2009.10.003

Sanz-Arigita, E. J., Schoonheim, M. M., Damoiseaux, J. S., Rombouts, S. A. R. B., Maris, E., Barkhof, F., Scheltens, P., & Stam, C. J. (2010). Loss of 'small-world'networks in Alzheimer's disease: graph analysis of FMRI resting-state functional connectivity. *PLOS ONE*, *5*(11), e13788.

Utianski, R. L., Caviness, J. N., van Straaten, E. C. W., Beach, T. G., Dugger, B. N., Shill, H. A., Driver-Dunckley, E. D., Sabbagh, M. N., Mehta, S., Adler, C. H., & Hentz, J. G. (2016). Graph theory network function in parkinson's disease assessed with electroencephalography. *Clinical Neurophysiology*, *127*(5), 2228–2236. https://doi.org/10.1016/J.CLINPH.2016.02.017

Watts, D. J., & Strogatz, S. H. (1998). Collective dynamics of 'small-world'networks. *Nature*, *393*(6684), 440.

Zalesky, A., Fornito, A., & Bullmore, E. T. (2010). Network-based statistic: identifying differences in brain networks. *Neuroimage*, *53*(4), 1197–1207. https://doi.org/10.1016/j.neuroimage.2010.06.041
